# Dissociation between physical reasoning and tool use in individuals with left hemisphere brain damage

**DOI:** 10.1101/2025.11.20.688863

**Authors:** Shuchen Liu, Seda Karakose-Akbiyik, Alfonso Caramazza, Filipa Dourado Sotero, Laurel J. Buxbaum, Aaron L. Wong

## Abstract

Many everyday tasks, from chopping vegetables to catching a ball, require understanding both how objects respond to physical forces and how to use them effectively. These capacities, tool use and physical reasoning, are often assumed to rely on shared cognitive and neural mechanisms. At some level, this correspondence is expected: using an object typically requires understanding its physical properties. However, both capacities are complex and multicomponential, so the relationship between them may vary across levels of representation and task demands. Here, we asked whether third-person physical reasoning about object dynamics can dissociate from tool use (i.e., performing a tool’s typical action, such as using a hammer to drive a nail) in individuals with left-hemisphere stroke. We tested 11 patients, five of whom showed impairments in tool use. Physical reasoning was assessed using a novel collection of tasks probing judgments about mass, velocity, and timing across static and dynamic scenes. Tool use was evaluated using a classic gesture-to-sight task, a pantomime-based measure in which participants are shown pictures of familiar tools and asked to demonstrate how each would be used. We identified an individual-level dissociation: patient I.A.* showed impairment in gesturing the use of objects despite preserved physical reasoning, often outperforming neurotypical controls. This pattern was complemented by patient N.P., who showed the reverse profile, with intact tool use gestures but difficulties in some physical reasoning tasks. These findings suggest that the ability to reason about the physical world and tool use can dissociate behaviorally and be independently disrupted by brain damage. This challenges the view that physical reasoning and tool use draw on the same underlying cognitive and neural mechanisms and suggests that at least some of their components are distinct.

## Introduction

Every moment we are awake, our brains interpret the physical structure and dynamics of the world to guide our actions. We predict where a moving vehicle will end up to decide whether it is safe to proceed. We lift, carry, and manipulate objects in ways that serve our goals. These everyday actions rely on an understanding of how objects move, how forces act on them, and how our actions influence these dynamics (i.e, physical reasoning about objects). How does the human mind support these abilities, and how are the cognitive processes that enable us to act in the world, from understanding physical dynamics to executing motor actions, related to one another?

Research in cognitive neuroscience has approached this question through multiple directions, primarily through studies of action planning and tool use (Lewis, 2006), and through investigations of physical reasoning (Fischer et al., 2016; Jack et al., 2013; Schwettmann et al., 2019). Yet the cognitive and neural mechanisms supporting these abilities are often studied separately. As a result, we lack a unified account of how physical reasoning and tool use are organized in the mind and brain, and under what conditions they may dissociate. To address this gap, we tested physical reasoning and tool use in individuals with left-hemisphere brain damage. Using a single-case series design, we tested the extent to which these abilities may dissociate in individual patients.

We define tool use as the ability to use familiar tools in conventional ways, such as knowing how to grasp and manipulate a hammer to drive a nail. Separately, we define physical reasoning as the ability to predict and interpret how objects react to external physical forces. These two capacities are often thought to rely on shared cognitive and neural mechanisms (Fischer & Mahon, 2021; Osiurak & Badets, 2016). At some level, a correspondence between physical reasoning and tool use is expected: using an object typically requires an understanding of its physical properties. For example, using a hammer to drive a nail requires knowledge not only of the hammer’s function but also of its mechanical properties, the forces involved in striking, the resistance of the nail, and the stability of the surface. Because humans interact with the physical world so naturally and fluently, the underlying processes are often intertwined, making them difficult to separate using behavioral paradigms alone. For this reason, researchers have often turned to other approaches: neuroimaging studies, which have typically examined the neural bases of tool use and physical reasoning in isolation rather than their interplay, and neuropsychological studies of patients, which test for possible dissociations in component processes.

### Cognitive neuroscience of physical reasoning and tool use

Previous research has identified a network of frontoparietal brain regions involved in planning, executing, or pantomiming actions, particularly those involving tools or manipulable objects (Gallivan & Culham, 2015). For instance, dorsal and ventral premotor cortex, superior parietal lobule, and anterior and posterior intraparietal sulcus show increased activity when participants manipulate objects like hammers, compared to control conditions such as grasping a neutral bar (Brandi et al., 2014). It has also been shown that these regions encode representations that can be used in support of actions, such as the direction of movement or the effectors that are involved (Gallivan et al., 2011).

Similar brain regions have also been implicated in third-person physical reasoning, the so-called *naive physics* network. For example, increased activity is seen in a network of frontoparietal regions when observers judge which direction a stack of blocks will fall, as opposed to reporting their superficial features like color (Fischer et al., 2016). These regions also carry information about the physical properties of objects when people observe but do not interact directly with them. For instance, the weight (Schwettmann et al., 2019), stability (Pramod et al., 2022), or trajectory (Karakose-Akbiyik et al., 2023, 2024) of objects in visually presented dynamic scenes can be decoded in these brain regions. Notably, brain regions involved in physical reasoning appear to overlap substantially with those involved in the planning and use of manipulable objects (Fischer et al., 2016; Fischer & Mahon, 2021; Osiurak et al., 2024).

Neural overlap has often been interpreted as evidence for shared neural and cognitive resources. In the original description of the so-called naive physics network, it was proposed that regions that are involved in physical reasoning overlap with regions “*involved in action planning and tool use, pointing to a close relationship between the cognitive and neural mechanisms involved in parsing the physical content of a scene and preparing an appropriate action*” (Fischer et al., 2016). More recently, this overlap has been interpreted more broadly as reflecting the role of these regions in understanding how actions both depend on and alter the physical dynamics of the world through a first-person physics engine (Fischer & Mahon, 2021). Some researchers have also proposed that tool use is fundamentally subserved by online technical reasoning about tool properties rather than by stored knowledge or motor routines, which may account for the apparent overlap between regions involved in tool use and those engaged in physical reasoning (Osiurak et al., 2024; Osiurak & Badets, 2016; for a discussion see Buxbaum, 2017).

Overlap between physical reasoning and tool use at some level is expected, as using an object typically requires an understanding of its physical properties. However, the extent to which they rely on shared mechanisms may be somewhat limited. For instance, the cognitive demands of familiar tool use may depend more heavily on learned motor routines, while using objects in novel situations might require engaging in physical reasoning more directly (Allen et al., 2019; Buxbaum & Randerath, 2018; Goldenberg & Spatt, 2009). Broader intuitions about how objects move or react to physical forces, such as expecting that a ball will accelerate as it rolls downhill, may draw on more abstract knowledge that has even less of a link to tool use. Thus, while some overlap in the underlying cognitive and neural systems is possible, the present study focuses on aspects of physical reasoning that may dissociate from tool use, particularly those involving third-person inference and prediction about object dynamics.

### Neuropsychological research on physical reasoning and tool use

Neuropsychological research also supports the role of frontoparietal regions (as well as the posterior temporal lobe) in tool use. Damage to these structures in the left hemisphere can result in limb apraxia, a disorder characterized in part by impaired use or pantomime of familiar or novel tools (Goldenberg & Spatt, 2009). For example, a person with apraxia might select an inappropriate tool for a task, such as a screwdriver instead of a spoon for eating, or use the correct tool in an ineffective manner, such as attempting to press a hammer into wood rather than striking with it (Randerath et al., 2009, 2010). These deficits can arise as a result of a disruption in the ability to properly represent and plan the movements associated with the tool (Buxbaum et al., 2000, 2005). While it has been suggested that these individuals are also impaired in the ability to reason about mechanical interactions between tools and objects (e.g., Goldenberg and Hagmann 1998), this has generally been tested only in the context of paradigms that require direct interactions with objects.

Although apraxia is frequently characterized by difficulties with familiar tool use, research on limb apraxia has long emphasized its heterogeneity. A common distinction is made between (1) stored knowledge about a tool’s conventional use, for example, knowing how to hold and manipulate scissors, and (2) the ability to reason about an object’s physical properties to guide action in novel situations, such as choosing a shoe rather than a rubber duck to hammer a nail (Daprati & Sirigu, 2006; Heilman et al., 1997). While the former draws on both physical principles and learned associations about typical body-object interactions, the latter can be accomplished based on current information about the environment and body (i.e., object affordances). Damage to certain left hemisphere regions can impair either or both abilities, but they can also dissociate: some patients struggle with familiar tool use yet retain an intact ability to manipulate objects in novel contexts, and vice versa (Goldenberg & Hagmann, 1998).

While prior work on limb apraxia and related disorders has revealed important dissociations between conventional and novel tool use, it has primarily focused on object selection and use tasks; situations that involve specifying how an object’s shape or mechanical properties afford a particular action, such as selecting the right protrusion for a tool to pull a ring (Goldenberg & Hagmann, 1998; Goldenberg & Spatt, 2009). These tasks differ from the kinds of paradigms typically used in studies of physical reasoning, which often require making predictions about how objects behave under external forces, often in non-action contexts (Balaban & Ullman, 2025; Battaglia et al., 2013). Much less is known about how brain damage affects this kind of third-person physical reasoning, and whether it can dissociate from the ability to use tools. To our knowledge, third-person physical reasoning has not been directly tested in individuals with brain damage.

### The current study

Here, we assessed whether third-person physical reasoning about object dynamics can dissociate from tool use in individuals with left-hemisphere stroke. Physical reasoning was assessed using a novel selection of tasks probing judgments about mass, velocity, and timing across static and dynamic scenes. Tool use was evaluated using a gesture-to-sight task, which is a commonly used sensitive measure of tool-use ability. In this task, participants were shown pictures of familiar tools and asked to pantomime how each would be used. Our goal was to examine whether these capacities can dissociate, as evidenced by some patients showing impairments in one but not the other. Observing such dissociations would offer insight into the relationship between physical reasoning and tool use, and suggest that they may depend, at least in part, on distinct neural and cognitive mechanisms.

## Methods

### Participants

Eleven stroke patients with chronic left-hemisphere brain damage (5 female; age range = 52–73 years, M = 61.09, SD = 6.96) participated in the study. Participants were included if they were 18-89 years old and had a stroke in the left hemisphere occurring at least 6 months prior to the study. Exclusion criteria included a history of additional neurological disorders, evidence of pre-existing dementia, history of a major psychiatric disorder, significant sensory deficits (e.g., confirmed blindness in both eyes, profound deafness that would impact the ability to hear task instructions), significant non-stroke-related movement impairment of the upper extremities (e.g., due to recent orthopedic injury), and recent history of alcoholism or substance abuse.

All participants provided informed consent and were compensated for their participation. The study was approved by the Institutional Review Boards of Harvard University and Thomas Jefferson University. Each participant completed a general neuropsychological screening, a tool use assessment, and a set of physical reasoning tasks. Structural T1-weighted MRI scans were acquired to generate lesion maps, which were drawn either manually or using an automated algorithm, and subsequently validated by a neurologist who was blind to the study goals (see Thibault et al., 2025 for more details on the analysis of the structural scans). The neuropsychological and tool use data were collected as part of a larger research effort at Thomas Jefferson University, which also included 15 age-matched neurotypical individuals who served as controls. The physical reasoning tasks were developed specifically for this study, and data from age-matched neurotypical controls were collected for comparison (see *Physical Reasoning Tasks* for details on control participants).

We adopted a broad inclusion criterion, chronic left-hemisphere stroke, since deficits in tool use and understanding mechanical properties of objects are much more commonly associated with damage to the left than right hemisphere (Buxbaum & Randerath, 2018; Goldenberg & Hagmann, 1998). Given the broad inclusion criterion, we did not expect to observe clear dissociations in every case as extensive brain damage can result in deficits in multiple functions. Moreover, because third-person physical reasoning had not previously been tested in individuals with brain damage, a broad initial approach allowed for an exploratory investigation. Upon analyzing individual response patterns, we identified two patients, hereafter referred to as I.A. and N.P. (fictitious initials with no relation to their real names), who together exhibited a dissociation between tool use (praxis) and physical reasoning. The remainder of the paper focuses on their behavioral profiles; data from the other patients are included in the Appendix.

**Patient I.A.** is a right-handed man, aged 61, with 16 years of education. He was able to speak and communicate effectively and showed near-ceiling performance on a standardized language screening (*Comprehension Subtest of the Western Aphasia Battery* = 9.95/10). His neurological status at the time of testing was mildly impaired (*NIH Stroke Scale* = 7). Structural neuroimaging revealed lesions in the left dorsal central sulcus, ventral to dorsal precentral sulcus, and dorsal premotor cortex (see Fig. 1). Lesion-based disconnection mapping using the BCBtoolkit (Foulon et al. 2018) revealed extensive disconnections in white matter pathways including the superior longitudinal fasciculus, frontal aslant tract, fronto-striatal fibers, cingulum, and inferior frontal regions (see Appendix C).

**Figure 1.**
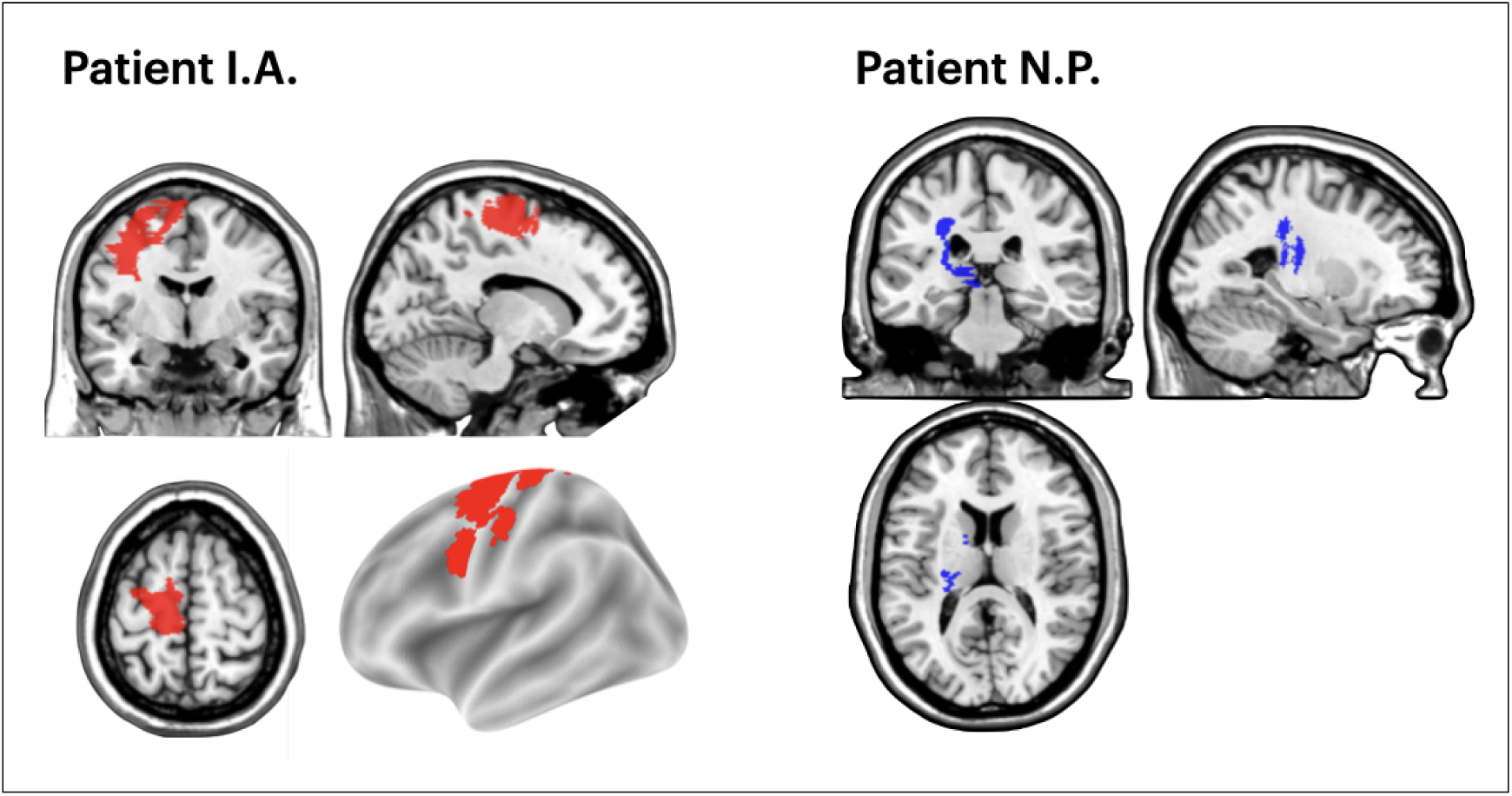
Lesion profiles of patient I.A. and N.P. Patient I.A. had lesions involving the left dorsal central sulcus, the region ventral to the dorsal precentral sulcus, and the dorsal premotor cortex. Patient N.P. exhibited a deep lesion involving the left thalamus and adjacent white matter, including the anterior limb of the internal capsule and the corona radiata with a largely spared cerebral cortex.

**Patient N.P.** is a 64-year-old right-handed woman with 12 years of education. She showed intact language abilities on a standardized screening (*Comprehension Subtest of the Western Aphasia Battery* = 10/10) and had mild neurological impairment at the time of testing (*NIH Stroke Scale* = 5). Structural neuroimaging revealed white matter lesions in the left hemisphere, with the cerebral cortex largely spared (see Fig. 1). Lesion-based disconnection mapping using the BCBtoolkit revealed disconnections within dorsal fronto-parietal pathways, including the superior longitudinal fasciculi, which links the superior parietal lobule to dorsal premotor regions (see Appendix C).

### Assessment of praxis ability

#### Gesture-to-Sight Task

Praxis was assessed using the gesture-to-sight task, a well-established measure of tool-use (Buxbaum et al., 2000, 2005). Participants were asked to pantomime the use of familiar tools with their left hand as all had left-hemisphere lesions that could compromise motor control of the contralateral (right) hand, and limb apraxia can still be observed on the ipsilesional (left) side.

Tool use pantomime is tightly linked to actual tool use in terms of performance (Belanger et al., 1994; Hermsdörfer et al., 2013), kinematics (Clark et al., 1994; Poizner et al., 1995), and involved brain regions (Osiurak et al., 2021). Overall, tool use pantomime is widely recognized as a sensitive method for assessing tool use in brain damage (Goldenberg, 1996; Randerath et al., 2011). Importantly, unlike real tool use, pantomiming removes structural and tactile feedback that can guide grasp configuration and movement execution, thereby providing a more direct index of stored action knowledge and motor planning (Baumard et al., 2014). Thus, in the current study, pantomiming was used as a proxy for tool use.

Performance was evaluated across four dimensions: content, hand posture, arm posture and trajectory, and amplitude and timing. This coding scheme has been extensively validated and applied in multiple studies from our laboratory to assess tool-use ability and characterize praxis errors, demonstrating strong reliability and sensitivity to behavioral performance. Trained coders rated responses using a comprehensive error classification system (Buxbaum et al., 2005), and inter-rater reliability across coders was high (Cohen’s Kappa > 0.85; Altman, 1999).

Errors were classified as follows: Content errors reflect semantically related actions; hand posture errors involve a hand configuration or hand movement; arm posture and trajectory errors concern the positioning and movement of the arm relative to the body; amplitude and timing errors reflect the size of the gesture (e.g. overly small or exaggerated) or grossly improper number of repetitions of the movement. For example, when pantomiming the use of scissors, a content error might involve a hammering motion, a hand posture error might involve using a closed fist, and an arm posture error could involve holding the arm rigidly instead of moving it forward. Amplitude and timing errors in this context might involve making overly large movements or failing to show the expected repetitive snipping motion. For each trial, a dimension received a score of 0 if an error was identified, or a 1 otherwise. If a content error was identified for that trial, the other errors were not scored.

An average was computed across the three non-content dimensions to obtain a praxis score for each individual, with higher scores indicating better performance. These scores were compared to data from neurotypical controls (*N* = 15, *M_control_age_* = 65.73, *SD_control_age_* = 10.28, 8 females, *M_control_gesture_sight_* = 0.94, *SD_control_gesture_sight_* = 0.04). For this and other tasks, to evaluate whether a patient’s performance was significantly below that of controls, we used the Crawford & Howell (1998) modified t-test, which is specifically designed for comparing a single case to a small control sample (Crawford & Howell, 1998). In addition to the *t* statistic, we also report *z_cc_*, a standardized effect size measure associated with the modified *t*-test (Crawford et al., 2010). Patients who scored below the control range in the gesture to sight task were classified as apraxic.

Patient I.A. scored 0.74, significantly below controls (*t*(14) = −5.81, *p* < .001, *z_cc_* = −6.00), and was classified as having apraxia. He did not make any content errors. I.A.’s errors were primarily in hand posture and amplitude / timing. For example, while demonstrating how to use a wrench, he did not adopt the proper closed-hand grip and displayed an insufficient amplitude of wrist motion, lacking the coordinated arm movement required to complete the turning action.

Patient N.P. scored 0.89, within the control range (*t*(14) = −1.50, *p* = 0.16, *z_cc_* = −1.55), and was classified as not having apraxia. Out of 34 trials, patient N.P. made two content errors: in one trial, she pantomimed applying deodorant to her face, and in another, she used a match as if it were a makeup implement. These trials were not analyzed further. N.P. performed well overall, although occasional hand posture errors were observed. For instance, in showing how to use a screwdriver, she used a pinch grip instead of a power grasp.

### Physical Reasoning Tasks

#### Stimuli and Apparatus

To investigate physical reasoning abilities, we developed a selection of tasks that required participants to make judgments about various physical properties of objects (e.g. mass, velocity) presented through dynamic or static scenes. Most tasks involved dynamic video stimuli created using the open-source animation software Blender. Three tasks (Weightlifting, Water, and Spring) used filmed or photographed real-world interactions instead. Each task began with a practice phase, which could be repeated at the participant’s request, or at the experimenter’s discretion if the participant needed more time to understand the task. Participants did not receive feedback for their practice trials or throughout the experiment.

Several tasks were adapted from previous paradigms used to target physical reasoning (e.g., Mitko et al., 2024). However, existing behavioral paradigms were typically developed to elicit a wide range of performance among neurotypical individuals, sometimes including tasks that even neurotypical individuals found challenging. Such designs are not optimal for assessing impairments in individuals with neurological impairment. To address this, we developed tasks that were relatively easy, ensuring that neurotypical participants could reliably understand and complete them. This approach made any deficits observed in the patient group easier to interpret. During task development, we benchmarked task performance in neurotypical control participants to ensure high (near ceiling-level) accuracy. Once finalized, the full battery was administered to age-matched neurotypical participants to establish reference performance levels (sample sizes = 9-15, depending on the task; see below). Control data were collected both in person and online via the Prolific platform.

#### General overview

We characterized our tasks broadly into three categories:

1. **Type A: Dynamic Mass Inference**: Tasks in which participants infer object mass from collision dynamics (see Fig. 2).
2. **Type B: Control Tasks**: Tasks designed to assess physical reasoning using alternative cues, such as human motion, static scenes, or indirect physical consequences (e.g., water splash, spring extension). These tasks functioned both to validate participants’ basic understanding of physical concepts in cases of impaired performance on dynamic tasks, and as assessments of perceptual components (e.g., velocity estimation) that are required for dynamic mass inference (see Fig. 3).
3. **Type C: Timing Judgments**: Tasks designed to assess participants’ ability to make inferences and predictions about timing based on dynamic interactions. These tasks also served as controls to determine whether individuals who failed mass inference tasks could nonetheless succeed on timing-based judgments (see Fig. 4). Note that predicting the timing of dynamic events and inferring objects’ physical properties are often grouped together under the umbrella of physical reasoning (Ullman, et al., 2017; Mitko, et al., 2024). We suggest, however, that these abilities may dissociate. Judgments about velocity and event timing can often be made by extrapolating an object’s trajectory without reference to its material properties, whereas inferring properties such as mass from collisions typically requires integrating multiple physical features, including force, resistance, and support relations. Given these considerations, we included timing-based tasks to assess whether participants who struggled with mass inference could nonetheless succeed on timing judgments.

**Figure 2.**
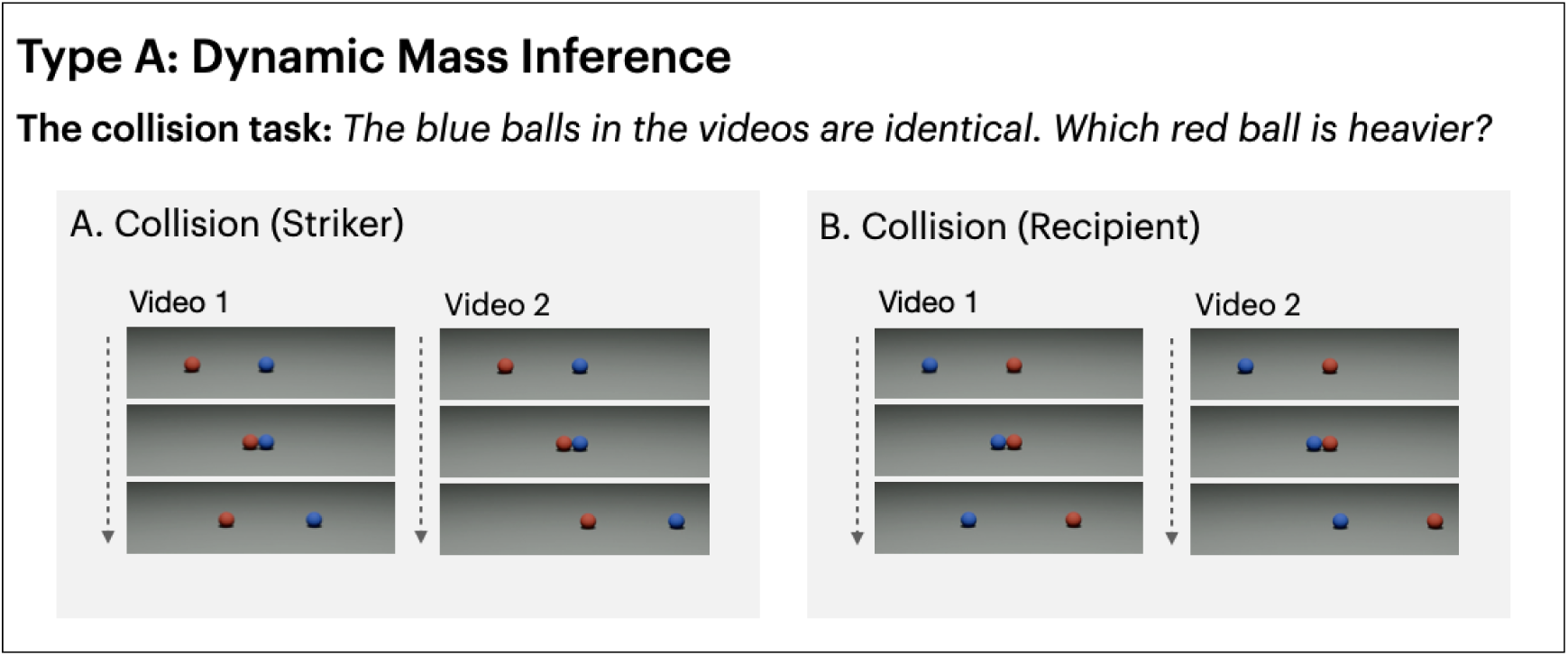
Examples of tasks in Type A (dynamic mass inference). (A) Collision task (striker version): on each trial, participants saw two videos of a red ball striking a blue ball. (B) Collision task (recipient version): on each trial, participants saw two videos of a blue ball striking a red ball. In both versions, the weight of the blue balls were the same between the videos. Participants compared the weight of the two red balls and judged which video had the heavier red ball.

**Figure 3.**
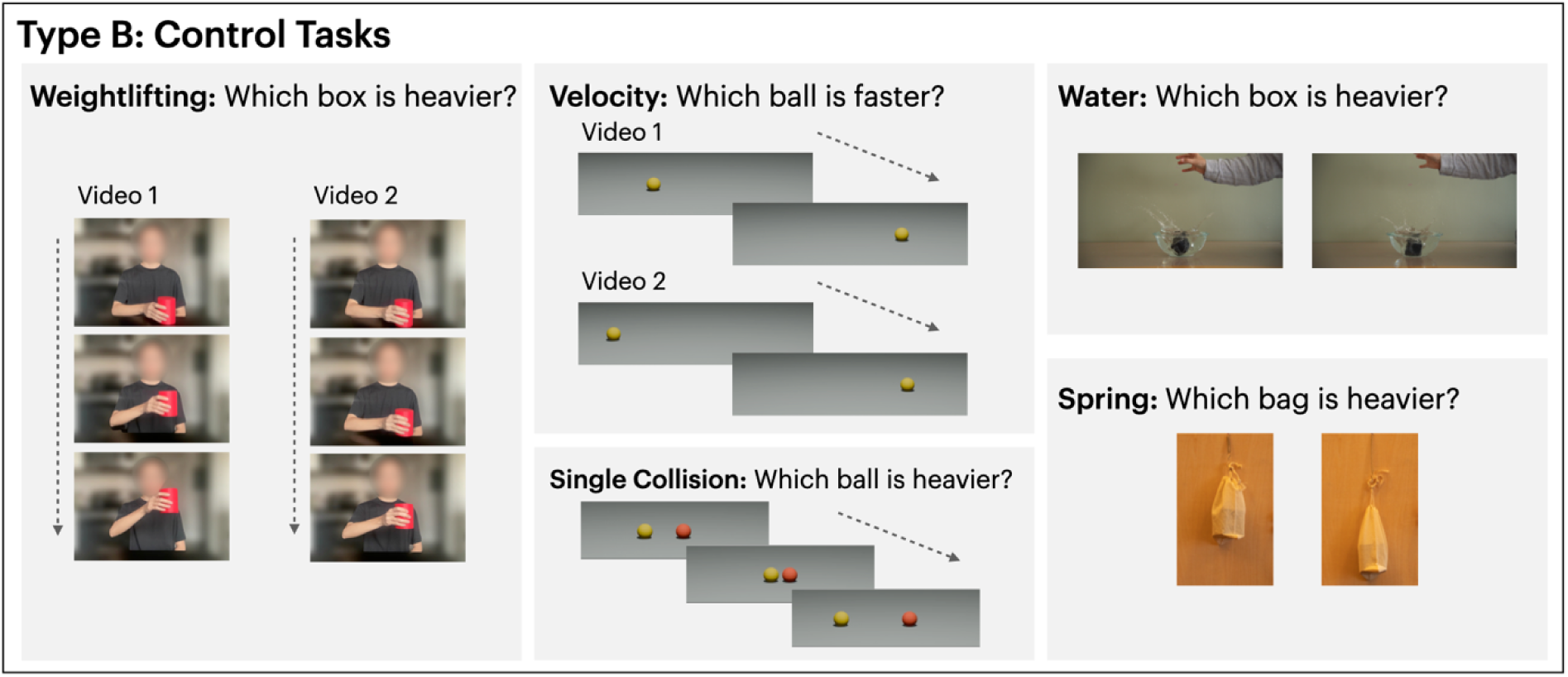
Examples of tasks in Type B (control tasks). Weightlifting task: on each trial, participants watched two videos of the same person lifting a container. The two videos were played in sequence. Participants judged which container was heavier. The paradigm was adapted from Mitko et al. (2024). For illustration purposes, one of the authors recreated the stimuli by filming themselves, as shown in this figure. The actual stimuli used in this task were a subset from of the stimuli from Mitko et al. (2024). Velocity task: on each trial, participants watched two videos of a ball rolling across a flat surface. Participants judged which ball was faster. Water task: on each trial, participants viewed two images capturing a box being dropped into a bowl of water. The boxes were visually identical but contained weights. Participants judged which box was heavier. Spring task: on each trial, participants viewed two images capturing a box hanging from a spring. The boxes were visually identical but contained weights. Participants judged which box was heavier.

**Figure 4.**
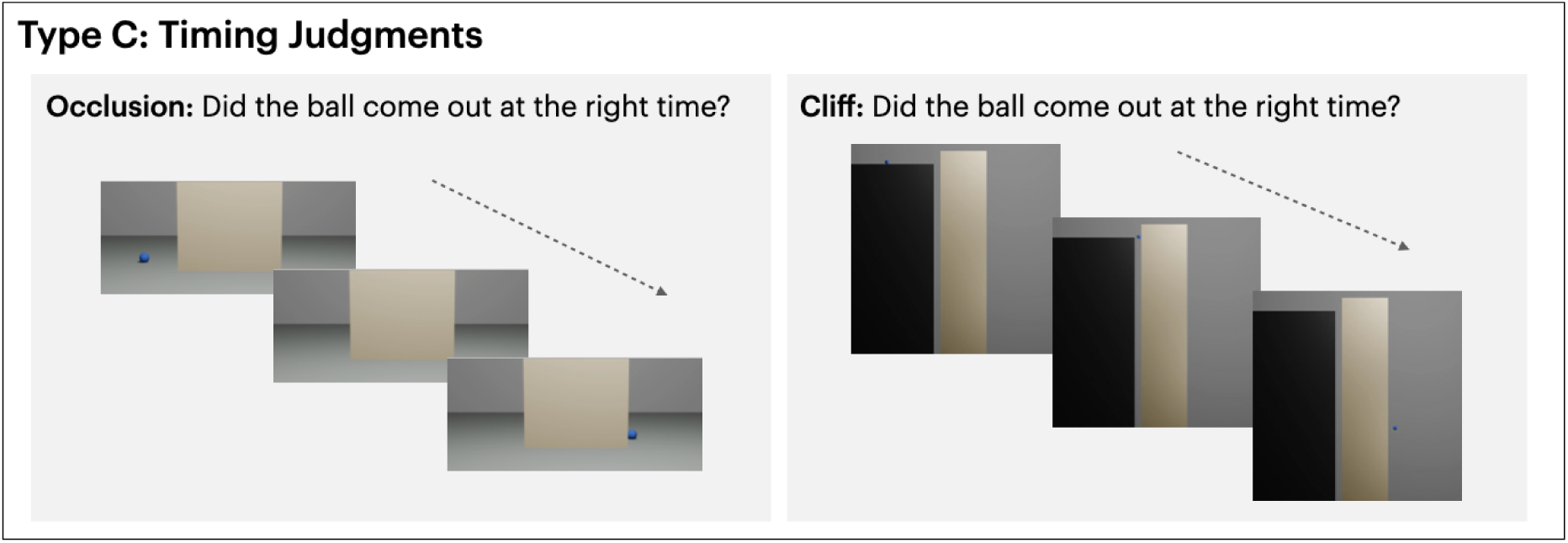
Examples of tasks in Type C (timing judgements). Occlusion task: on each trial, participants watched a video of a ball rolling at a constant speed, going behind an occluder and coming out the other side after some time. Participants judged whether the ball came out of the occluder at the correct time. Cliff task: on each trial, participants watched a video where a ball rolled at a constant speed off a high platform, started falling and went behind an occluder next to the platform. It came out of the occluder from the other side after some time. Participants judged whether the ball came out of the occluder at the correct time.

Even though neurotypical control participants often reached high levels of accuracy across our tasks, some paradigms included trials that were more challenging, leading to systematic error patterns in certain neurotypical individuals. To characterize these patterns, we evaluated whether control participants exhibited consistent errors for specific trial types. For example, in the collision tasks, some controls systematically misjudged scenarios where the striking ball continues moving after the collision in one video but stopped in another. We did not want to penalize stroke participants for errors in such cases, given that these scenarios may reflect common biases or heuristics in physical reasoning (Caramazza et al., 1981; Gilden & Proffitt, 1989; Kaiser et al., 1985; Mitko & Fischer, 2023) rather than a deficit not found in neurotypical subjects.

In cases where such biases were observed, we adjusted task scores by excluding these trial types for all participants (both patients and controls) to ensure fair comparisons. This adjustment allowed us to assess whether patients’ errors were qualitatively similar to those of controls and to avoid underestimating patient or control performance based on error patterns that are common in neurotypical individuals. These adjustments are described in more detail in the relevant task sections below. For all tasks, we used the Crawford & Howell (1998) modified t-test to compare each patient’s performance to that of the control group to determine whether a patient’s performance was statistically atypical relative to age-matched neurotypical participants. In addition to the *t* statistic, we also report *z_cc_*, a standardized effect size measure associated with the modified *t*-test (Crawford et al., 2010).

### Type A: Dynamic Mass Inference

#### Collision Task

This task probed participants’ ability to infer the relative mass of objects based on dynamic interactions; specifically, the outcome of collisions between two balls. It was adapted from previous paradigms involving colliding objects (Mitko & Fischer, 2023; Runeson et al., 2000; Todd & Warren, 1982). On each trial, participants viewed two vertically aligned videos played simultaneously, each depicting a red ball colliding with a blue ball. The collisions occurred at the center of the screen. The balls were visually identical in appearance apart from their color.

Each video began with one ball stationary and the other moving at a constant speed. There were two versions of the task: *red ball as striker* and *red ball as recipient*. In the striker version (Fig. 2A), the red ball entered the screen and collided with the stationary blue ball. In the recipient version (Fig. 2B), the red ball remained stationary and was struck by an incoming blue ball. Participants were told that the blue ball always had the same mass, and that one of the red balls (top or bottom video) was heavier than the other red ball. Their task was to report which red ball appeared heavier by pressing a button. Each video pair played twice per trial, and the two versions were presented in counterbalanced A–B–B–A blocks (24 trials per block, 48 trials per version). The animations followed idealized Newtonian physical laws with no energy loss or surface friction. The blue ball’s weight was fixed at 10 units. The red ball’s weight was set to 5, 10, or 20 units, allowing for weight comparisons of 20–5, 20–10, and 10–5 across videos.

Creating the two versions of the collision task was important, as the post-collision behaviors of the recipient and the striker are differently affected by the relative mass between the striker and the recipient. For instance, a recipient ball always moves forward when struck, and the lighter it is, the faster it is. Conversely, a striker’s motion post-collision depends on whether it is heavier or lighter than the recipient: it continues forward if heavier, rebounds if lighter, and remains stationary if equal. Thus, simple heuristics (e.g., “the heavier ball moves less”) might work in one version but fail in the other. Successfully performing this task requires a flexible reasoning system that can adjust to the causal structure of each scenario, rather than relying on fixed visual cues or one-size-fits-all rules.

#### Easy Collision Task

This task used the same structure as the standard collision task but used more extreme mass differences to reduce difficulty. It served as a control to determine whether participants who struggled with the original version could succeed when mass contrasts were more salient. As before, participants compared two videos showing red and blue balls colliding at the center of the screen and judged which red ball appeared heavier. The blue ball’s weight was fixed at 30 units, while the red ball varied among 1, 6, 20, 36, 75, and 600 units. Unlike the standard task, no condition included equal masses between the red and blue balls. In half of the trials, both striker balls were heavier than the recipient and moved forward after the collision; in the other half, both striker balls were lighter than the recipient and bounced back after the collision.

### Type B: Control tasks

Control tasks were designed to assess participants’ understanding of the foundational components of physics that are required to perform well in the collision tasks. Specifically, we hoped to: (1) confirm that participants had an intact general sense of mass, (2) confirm that participants could distinguish different velocities, an essential cue to solving the collision tasks in Type A, (3) test whether participants could make inferences of mass in alternative non-collision contexts, and (4) determine whether participants could succeed in easier versions of the collision tasks that can be solved based on simple heuristics. Together, these tasks provided a framework for interpreting poor performance on the more complex collision trials.

#### Single Collision Task

In this simplified version of the collision task, participants viewed a single video per trial showing a red and a yellow ball colliding. The balls were visually identical except for color. Participants indicated which ball (left or right) they believed was heavier via button press. On half the trials, the red ball acted as the striker; on the other half, the yellow ball did. In half of the trials, the recipient ball’s mass was fixed at 1 unit, while the striker ball’s mass varied across 4, 6, 8, or 10 units. In the other half, the striker ball’s mass was fixed at 1 unit, while the recipient ball’s mass varied across 4, 6, 8, or 10 units. Each participant completed 4 practice trials followed by 16 test trials.

The mass pairings were selected such that the heavier ball always moved more slowly after the collision regardless of whether it was the striker or the recipient. As a result, the simple heuristic “the slower-moving ball is heavier” would lead to perfect accuracy in this task. In contrast, this same heuristic is misleading in the standard collision task, where post-collision speed depends on the specific role (striker vs. recipient) and relative mass of the objects. Therefore, the Single Collision Task served as a control condition, where participants could succeed using a straightforward perceptual rule, even if they lacked more flexible or context-sensitive physical reasoning.

#### Velocity Task

This task assessed participants’ ability to judge speed differences, a critical skill underlying mass inference in the collision paradigms. In collision events, speed differences are key cues to relative mass: after being struck with the same force, a heavier ball moves more slowly than a lighter ball, assuming other factors (e.g., friction, air resistance) are held constant. Thus, this task served as a reference point to interpret failures on collision tasks to see whether they stemmed from difficulties in physical reasoning or could be explained by basic difficulties in comparing object velocities.

In each trial, participants saw two vertically aligned videos, each showing a ball entering the screen, rolling at a constant speed, and exiting. The two balls moved at different speeds, and participants indicated which one moved faster. The task included 4 practice trials and 48 test trials. To prevent participants from relying on simple heuristics based on entry or exit times rather than speed itself, entry times were offset across trials. In 25% of trials, the slower ball entered the screen first and exited before the faster ball caught up, ensuring that participants had to judge true speed rather than rely on timing-based shortcuts.

#### Water Task

This task assessed participants’ ability to infer object weight from indirect physical effects captured in static images. In each trial, participants viewed two photographs showing a visually identical box dropped into a bowl of water, captured at the peak of the resulting splash. Participants were asked to indicate which box was heavier. The difference in weight was created by placing unseen items inside the boxes. The heavier box weighed either 40g or 50g, and the lighter box weighed either 0g (an empty box) or 10g. Videos of the splashes were recorded, and screenshots were selected from the moment most predictive of weight differences. Each participant completed 4 practice trials and 32 test trials.

#### Spring Task

This task also used static images to probe participants’ intuitive understanding of weight, this time through the deformation of a spring. In each trial, participants viewed two photographs of the same bag suspended from a spring, with the only difference being the weight inside the bag. Participants were asked to select which bag appeared heavier. The spring and bag setup was held constant across trials, with weights hidden inside a box placed within the bag to minimize visual confounds. The weights varied from 0g to 500g across six levels. Each participant completed 4 practice trials and 60 test trials.

#### Weightlifting Task

This task assessed participants’ ability to judge the relative mass of objects based on human motion cues, rather than direct object-object interactions. We directly adapted the paradigm from Mitko et al. (2024), using a subset of their stimuli where an actor or actress is lifting visually identical containers of different weights. On each trial, participants viewed two videos (presented in an alternating ABAB sequence) and were asked to indicate which video showed the actor lifting a heavier container. In each trial, the two videos depict the same actor or actress; the heavier container weighed either 18 or 20 units, and the lighter container weighed either 1 or 3 units. Each participant completed 4 practice trials and 32 test trials

### Type C: Timing tasks

Inference of mass is only one aspect of reasoning about physical interactions. We often need to infer properties of a dynamic scene that are not inherent to a specific object, such as location, trajectory, and timing. To further explore the patients’ reasoning about other types of inferences about dynamic scenes, we designed two exploratory tasks probing participants’ ability to make predictions about the temporal aspect of a scene.

#### Occlusion Task

This task assessed participants’ ability to make temporal predictions based on velocity and distances. In each trial, a video showed a ball rolling at a constant speed across a flat surface and passing behind a vertical occluder at the center of the screen. The ball disappeared behind the occluder and re-emerged from the other side at a specific time point. Participants were asked to judge whether the ball reappeared at the *correct* time, assuming uninterrupted motion behind the occluder, or whether it appeared at the wrong time (too early or too late). Videos were rendered at 30 frames per second (FPS). In 50% of the trials, the timing of reappearance was physically accurate. In 25% of the trials, the ball reappeared early (23, 26, or 31 frames too soon), and in the other 25%, it reappeared late (23, 26, or 31 frames too late). Two different ball speeds were used to introduce variation. There were 8 practice trials and 72 test trials.

#### Cliff Timing Task

The cliff timing task was a more complex variant of the occlusion task. A ball rolled horizontally at a constant speed across a raised platform (the “cliff”), then dropped off the edge and disappeared behind a vertical occluder during its fall. Participants judged whether the ball reappeared at the correct time on the other side of the occluder. Crucially, the correct reappearance time was determined solely by the ball’s initial horizontal velocity. Assuming ideal physics, gravity would dictate the vertical position of the ball, while horizontal ball velocity would dictate its horizontal position when falling. Since the occluder was vertical and the same height as the cliff (i.e., the ball can’t fall “below” the occluder), the time of reappearance was dependent on the ball’s horizontal velocity relative to the width of the occluder, not gravity. Thus, participants should ignore irrelevant gravitational cues. Each video was presented at 30 frames per second. In 50% of the trials, the ball reappeared at the correct time; in 25%, it reappeared early (17 or 22 frames too soon); and in 25%, it reappeared late (28 or 36 frames too late). Two different ball speeds were used to introduce variation. There were 8 practice trials and 64 test trials

## Results

For each task, we collected data from 9 to 15 age-matched neurotypical controls, either in-person or online via the Prolific platform. Among the 11 left-hemisphere stroke patients tested, no significant correlation was found between any physical reasoning task and praxis score (see Appendix B, Table 2). Furthermore, we identified two individuals, patients I.A. and N.P., who exhibited a dissociation between praxis and physical reasoning abilities. Patient I.A., classified as having apraxia, performed exceptionally well on the physical reasoning tasks, providing evidence for a dissociation between praxis and physical reasoning. This pattern was complemented by patient N.P., classified as not having apraxia, who showed notable difficulties in several of the physical reasoning tasks. Below, we present each patient’s performance alongside the control range for each task (see Appendix B Table 1 for control range). Data from the remaining patients appear in the Appendix A.

### Type A: Dynamic Mass Inference

#### Collision Task

This task assessed the participants’ ability to infer relative object weight from motion cues in collision events. They watched two vertically aligned videos depicting a red ball colliding with a blue ball. Across the two videos, the two blue balls were always the same weight, whereas the two red balls had different weights (see Methods for details). Participants judged which red ball was heavier. We tested two versions of this task: *red ball as striker*, where the red ball initiated the collision, and *red ball as recipient*, where the red ball was being struck. In both versions, the blue ball in the two videos always had the same weight (10 units) and the trials varied based on the red ball weight pairings in the two videos (20–10, 20–5, 10–5), which produced different post-collision movement patterns. We conducted a repeated-measure ANOVA across trial types on the control subjects to search for systematic error patterns

#### Red ball as recipient

In judging the weight of the recipient ball, control participants performed with a mean accuracy of 93.91% (N = 13, SD = 7.39%, range: 83.33% - 100.00%). Patient I.A. scored 100.00%; Patient N.P. scored 31.25%. There was no significant pattern of errors across trial types in controls (*F*(1.16, 13.88) = 2.33, *p* = .15, η^2^_g_ = 0.08), so no score adjustments were made. Patient I.A. scored within the control range (*t*(12) = 0.79, *p* = .44, *z_cc_* = 0.82), ranking above 55.75%. Patient N.P.’s score was significantly below the control range (*t*(12) = –8.17, *p* < .001, *z_cc_* = −8.48), placing her lower than 100% of controls (see Fig. 6).

#### Red ball as striker

In judging the weight of the striker ball, control participants performed with a mean accuracy of 80.93% (N = 13, SD = 22.37%, range: 37.50% - 100.00%). In this initial unadjusted version of the task, patient I.A. scored 100.00% while patient N.P. scored 52.08%. A repeated-measures ANOVA revealed that controls made significantly more errors on 20–10 trials (*F*(1.17, 14.01) = 6.65, *p* = .02, η^2^_g_ = 0.17). We therefore adjusted the scores by excluding these trials. After score adjustment, controls averaged 89.18% (SD = 17.80%), with a range of 50.00%–100.00%. Patient I.A.’s score remained at 100.00%, while Patient N.P.’s dropped slightly to 50.00%, suggesting her errors differed from the typical errors of the controls. Patient I.A.’s performance did not differ from controls (*t*(12) = 0.59, *p* = .57, *z_cc_* = 0.61), ranking above 43.11%. Patient N.P.’s performance was marginally below the control range (*t*(12) = –2.12, *p* = .055, *z_cc_* = −2.20), placing her below 94.46% of controls (see Fig. 6). Across the two conditions, Patient I.A., despite being apraxic, performed perfectly on both versions of the collision task. In contrast, Patient N.P., despite preserved abilities of tool use pantomime, demonstrated a notable deficit (see Fig. 5).

**Figure 5.**
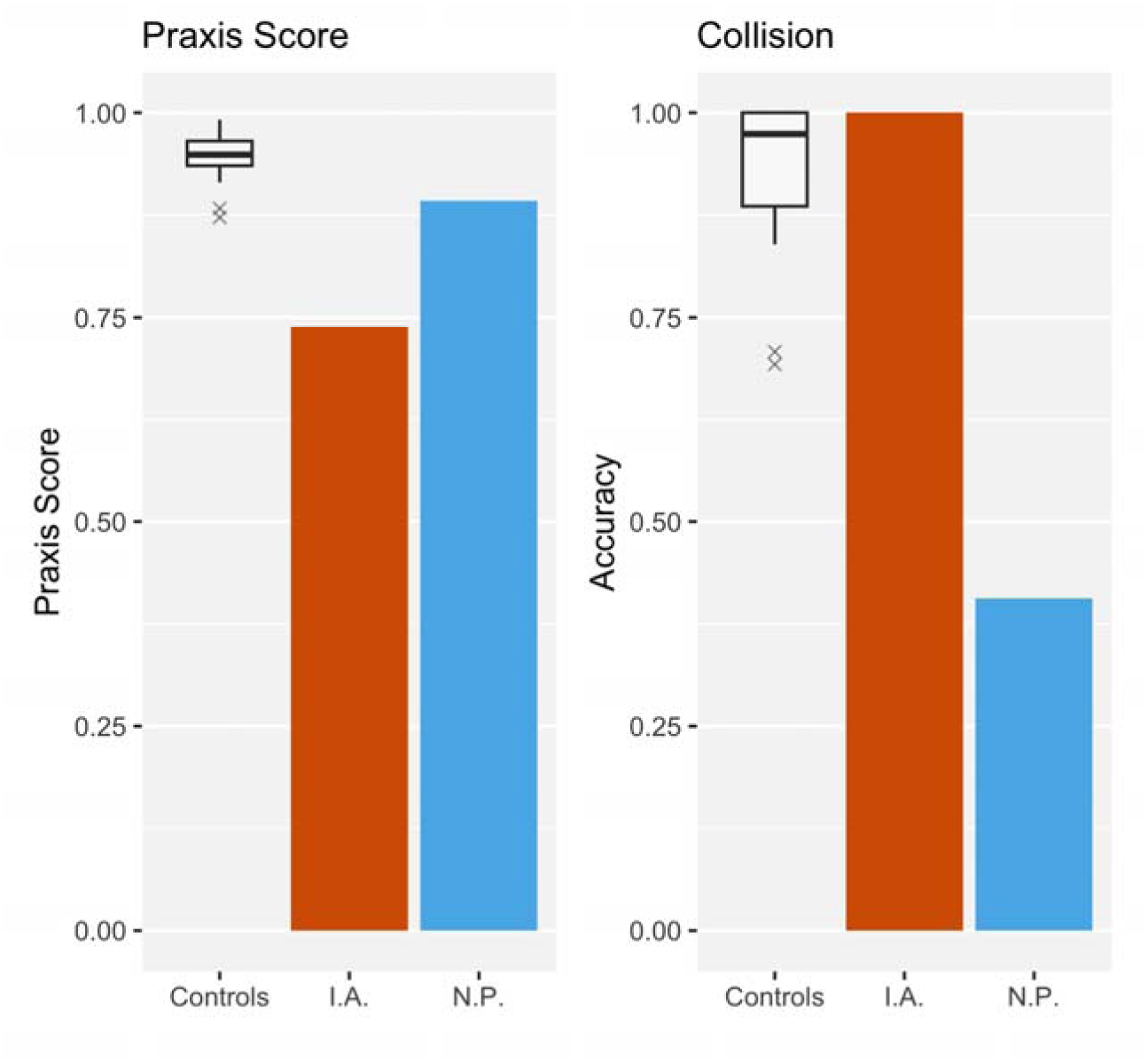
Comparison of praxis and judgment of mass in the collision task. The accuracy of the collision task reflects the average of the striker and recipient versions. Individual performance for I.A. (orange) and N.P. (blue) is shown alongside neurotypical controls, whose group distribution is displayed as a boxplot.

**Figure 6.**
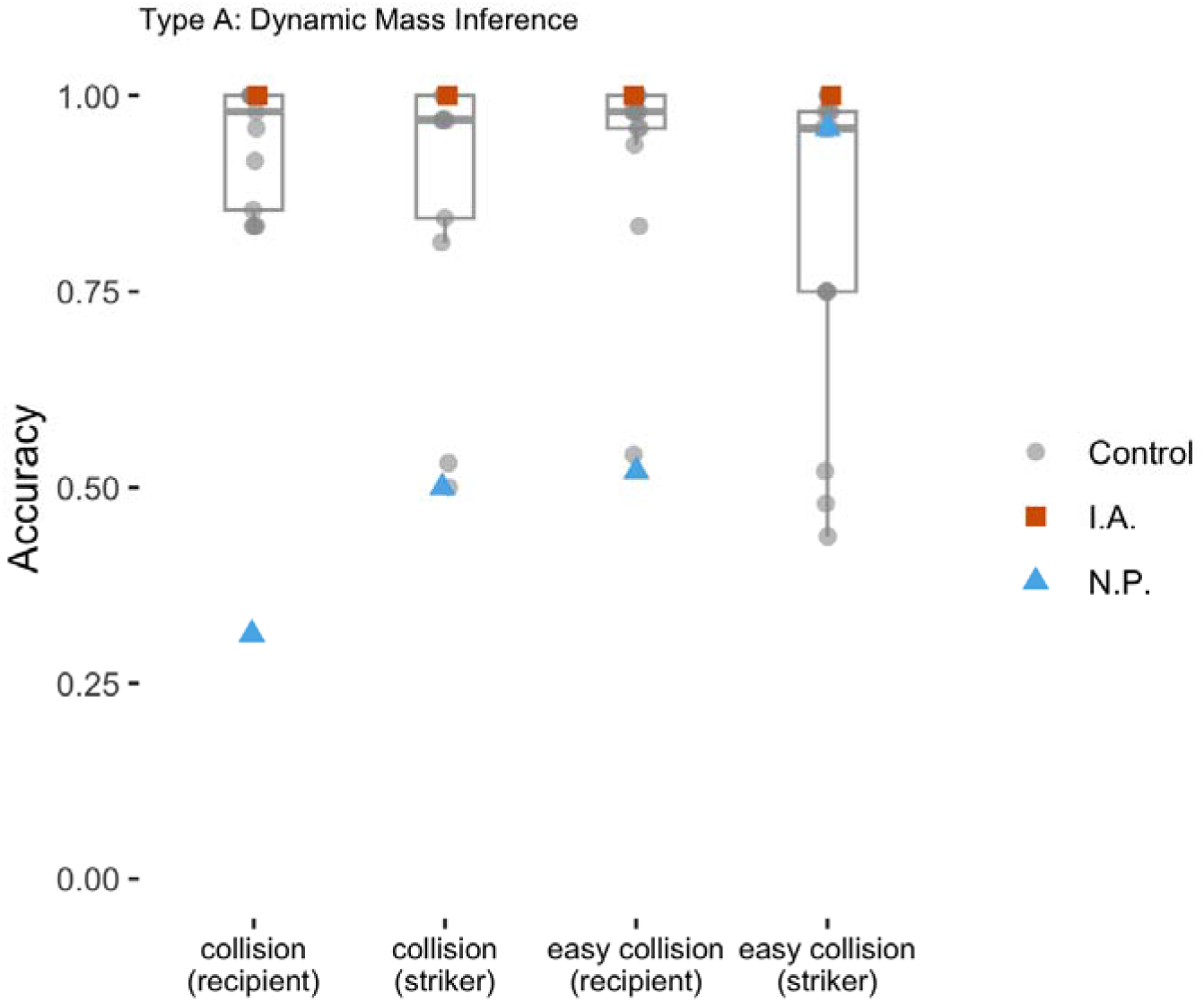
Accuracies for Type A - Dynamic Mass Inference tasks. The box plots showed the distribution of the control subjects. The individual dots showed the accuracy of individual subjects (gray circle: controls; red square: I.A.; blue triangle: N.P.)

#### Easy Collision Task

In the Collision task described above, some trials included a special case where the striker ball and the recipient ball had equal weight. In these cases, the striker ball came to a complete stop after impact, as dictated by the simulation’s idealized physics. Such perfectly elastic equal-mass collisions are rare in real life and may have been confusing for some participants. On the whole, poor performance on the Collision task might have stemmed from mass differences being too subtle or from participants being thrown off by the equal-weight trials. To address these possibilities and further assess the participants’ ability to infer relative mass from dynamic cues, we developed an easier version of the collision task. In this Easy Collision task, we removed equal-weight collisions and exaggerated mass differences between the two red balls in each trial to increase discriminability (ranging from 2.08:1 to 20:1). If participants’ difficulties with the original task were due to subtlety or confusion, we would expect them to perform reliably better in the Easy Collision task. Conversely, if they continued to struggle, this would suggest a more fundamental difficulty in understanding how physical laws govern object motion and interactions during collisions.

#### Red ball as recipient

Control participants showed an average accuracy of 94.31% (SD = 11.90%, range: 54.17% - 100.00%P). Patient I.A. again scored 100.00% while patient N.P. scored 52.08%. We found no significant error patterns across red ball weight combinations (*F*(2.61, 36.58) = 0.68, *p* = .55, η^2^_g_ = 0.01), thus, made no adjustments. Patient I.A.’s score was not significantly different from the control group (*t*(14) = 0.46, *p* = .65, *z_cc_* = 0.48), ranking above 34.97%. In contrast, Patient N.P.’s score was significantly lower than the control range (*t*(14) = –3.43, *p* = .004, *z_cc_* = −3.55), below 99.60% of controls. In this easy version of the collision task, only one control participant showed a similarly low overall accuracy to N.P.; however, their performance remained consistently near chance across both blocks (50% and 58%, respectively), suggesting that they were responding at random rather than performing the task as intended. In contrast, while N.P.’s average accuracy was 52.08%, she performed 95.83% in the first block and only 8.33% in the second. This sharp discrepancy is unlikely to reflect fatigue or lapses in attention. Rather, her reported strategy suggests the use of an internally consistent but incorrect rule. When asked about her approach in the second block, where she consistently failed, she explained that she chose the option in which “the red ball moved faster”. This strategy aligns perfectly with her responses but is systematically wrong: in this version of the collision task, where the red ball is the recipient, a slower recipient ball always indicates greater mass. Overall, N.P.’s performance tended to fluctuate depending on the arbitrary strategy she adopted at a given time. Thus, while her average score in this task resembled that of the control participant who also failed, the underlying pattern was qualitatively distinct, reflecting a rule-based but misguided reasoning rather than random guessing or inattention to the task.

#### Red ball as striker

Control participants achieved an average accuracy of 82.44% (N = 14, SD = 21.08%, range: 43.75% to 100.00%). Patient I.A. scored 100.00% while patient N.P. scored 95.83%. We evaluated trial difficulty based on red ball weight pairs (600–75, 600–36, 75–36, 20–6, 20–1, 6–1). A repeated-measure ANOVA found no significant error pattern across trial types (*F*(1.43, 18.64) = 2.33, *p* = .14, η^2^_g_ = 0.07), so no adjustments were applied.

Patient I.A.’s score was not significantly different from the control group (*t*(13) = 0.80, *p* = .44, *z_cc_* = 0.83), ranking above 56.45%. Patient N.P.’s score was also within the control range (*t*(13) = 0.61, *p* = .55, *z_cc_* = 0.64), ranking above 45.00%. These results indicate that both patients performed well on this simplified version in which the red ball was the striker, suggesting that N.P. maintained a relatively consistent, successful strategy in these blocks.

### Interim Summary & Discussion: Dynamic Mass Inference

Together, I.A. and N.P. exhibited a dissociation between praxis and judging the weight of interacting objects (see Fig. 5). Across these collision-based tasks, Patient I.A. consistently achieved perfect scores despite showing impairments in tool use. This suggests that the kind of physical reasoning required in these tasks (i.e., inferring object mass from kinematics) can remain intact in the presence of apraxia, and the ability to use tools and an intact physical reasoning capacity can dissociate. This is further supported by data from other patients, which show no clear relationship between praxis ability and accuracy on the collision-based tasks (see scatterplots of task performance vs. praxis scores in Appendix A).

Patient N.P., despite intact abilities related to tool use, showed a highly inconsistent pattern on the dynamic mass inference task. She performed well in the *striker version* of the easy collision task. However, she performed near chance in both versions of the original collision task and exhibited a variable performance across blocks in the *recipient version* of the easy task. This inconsistency suggests that her reasoning about physical interactions might be unreliable or dependent on superficial strategies rather than a stable internal model of physical principles.

At first glance, patient N.P.’s performance on the collision tasks appears impaired. However, it is important not to draw conclusions too quickly without ruling out alternative explanations. Because success on the collision tasks may depend on several contributing processes, we assessed a broader set of potential factors that could impact performance. One possibility is a general impairment in judging velocity, a critical perceptual cue for inferring mass from dynamic events Another possibility is a broader deficit in the intuitive understanding of mass. That is, rather than struggling with the specific cues used in the collision paradigms, an individual may have difficulty grasping the concept of mass itself, what it means for one object to be heavier than another, and how mass relates to physical interactions in general. Finally, it could be possible to have an intact understanding of mass and velocity but still fail to derive mass based on motion cues. To explore these possibilities and better interpret performance on the collision tasks, we developed a set of control tasks that target these foundational abilities more directly and test whether failure in the collision tasks can be explained by them.

### Type B: Control Tasks

#### Single Collision Task

In this task, participants judged relative weight in a single video of two colliding balls. Prior research has discovered a *striking bias*, a tendency to overestimate the weight of the striking ball when weight differences are subtle (Mitko & Fischer, 2023). We mitigated this bias by using trials with large weight differences. Ten controls achieved an average accuracy of 95.62% (*SD* = 5.76%), ranging from 81.25% to 100.00%. Patient I.A. scored 100.00%; Patient N.P. scored 89.58%. We found no systematic error pattern across weight combinations (*F*(2.05, 18.49) = 2.80, *p* = .09, η^2^_g_ = 0.14), so no score adjustments were applied. Patient I.A.’s score was comparable to the control group (*t*(9) = 0.72, *p* = .49, *z_cc_* = 0.76), ranking above 51.23%. Patient N.P. also fell within the control range (*t*(9) = –1.00, *p* = .34, *z_cc_* = −1.05), ranking above 34.37% (see Fig. 7). Thus, in a simplified setting with drastic weight differences and a single scenario to track, N.P. was able to distinguish heavy from light objects based on dynamic cues. Note that in this task, a simple heuristic, ‘*the heavier ball will always move slower*’, is sufficient to achieve perfect performance. By contrast, in the collision tasks described under Type A, successful performance requires flexibly adjusting this rule, since the relationship between weight and speed differs for the striker and recipient of a collision. Overall, N.P. was able to infer weight when the mapping between mass and motion was straightforward, but as shown above, struggled when correct performance required flexible, context-dependent reasoning.

**Figure 7.**
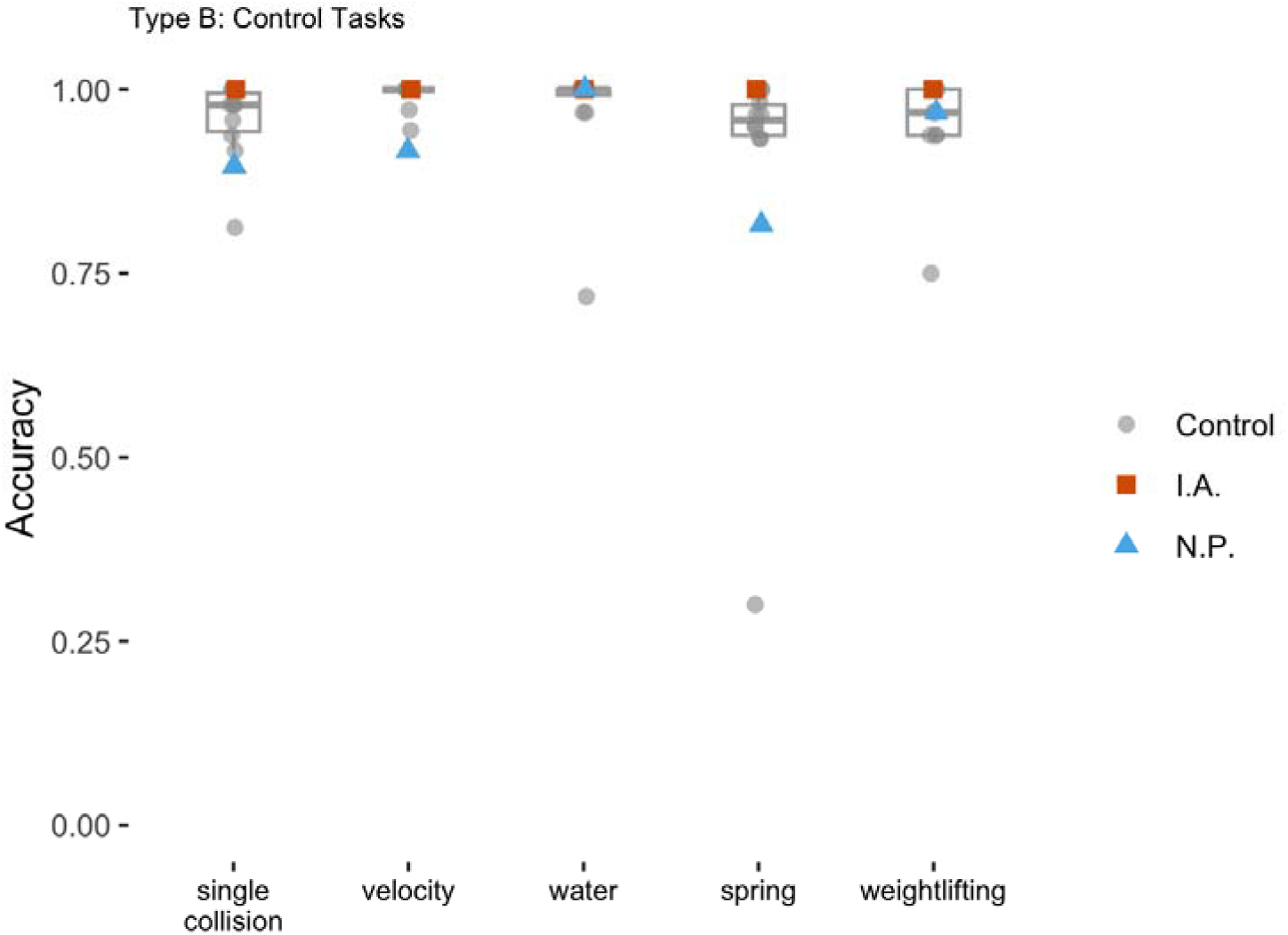
Accuracies for Type B - Control tasks. The box plots showed the distribution of the control subjects. The individual dots showed the accuracy of individual subjects (gray circle: controls; red square: I.A.; blue triangle: N.P.)

#### Velocity Task

In the collision paradigms, speed is a critical basis for inferring the relative mass of the balls. Thus, we tested if participants can judge the speed of moving objects. Participants were asked to judge which of two moving balls was faster. Control participants performed with a mean accuracy of 93.11% (*SD* = 8.35%, range: 75.00% - 100.00%). Patient I.A. scored 100.00%, and Patient N.P. scored 70.83%.

The velocity task included trials where the velocity difference between the two videos cannot be told from simple heuristics, leading to systematic errors for some trials even in neurotypical individuals. For instance, we had two types of trials: “catch-up” or “non-catch-up.” In non-catch-up trials, the slower ball entered and exited the screen before the faster ball could catch up, making judgments more difficult. Control participants made significantly more errors on non-catch-up trials (*t*(25) = –2.76, *p* = .01, *d* = 0.72). Thus, we also explored Patient N.P. and I.A’s performance in the adjusted scores by excluding these trials.

After adjustment, the control group’s mean accuracy rose to 99.36% (*SD* = 1.66%), ranging from 94.44% to 100.00%. Patient I.A. remained at 100.00%, while Patient N.P.’s score improved to 91.67%, indicating that many of their original errors mirrored those of controls (see Fig. 7). Modified t-tests showed that Patient I.A.’s adjusted score did not differ significantly from the control group (*t*(12) = 0.37, *p* = .72, *z_cc_* = 0.39), ranking above 28.3% of controls. Patient N.P.’s adjusted score was significantly lower than that of controls (*t*(12) = –4.45, *p* < .001, *z_cc_* = −4.62), below 97.55% of the control population. Even though Patient N.P. performed worse than most controls, she remained reasonably accurate at judging the velocity of the moving objects, especially for easier trials. Thus, her difficulty in the collision tasks described under Type A are unlikely to be attributable simply to an inability to judge velocity.

#### Water Task

In this task, participants viewed two images showing boxes dropped into water, with splash height reflecting weight. Control participants achieved an average accuracy of 97.14% (*N* = 12, *SD* = 8.05%, range: 71.88% −100.00%). Patient I.A. scored 100.00%; Patient N.P. scored 100% (see Fig. 7). We found no systematic error pattern across weight combinations (*F*(3, 33) = 1.688, *p* = .19, η^2^_g_ = 0.06), so no score adjustments were applied. Both scoring 100%, I.A.’s and N.P.’s performances were comparable to the control group (*t*(11) = 0.34, *p* = 0.74, *z_cc_* = 0.36), ranking above 26.13%.

#### Spring Task

Participants viewed images of a bag hanging from a spring, with weight inferred from spring extension. Control participants achieved an average accuracy of 89.83% (*N* = 10, *SD* = 21.16%, range: 30.00% - 100.00%) Patient I.A. scored 100.00%; Patient N.P. scored 81.67%. To assess systematic error patterns in controls, we conducted a repeated-measure ANOVA across weight combinations and found a significant effect of mass combination (*F*(14, 126) = 2.318, *p* = .007, η^2^_g_ = 0.06). However, post-hoc pairwise comparisons with Holm corrections did not reveal any significant difference among any pairs (all ps > 0.99), so no score adjustments were applied. Patient I.A.’s score was comparable to the control group (*t*(9) = 0.46, *p* = .66, *z_cc_* = 0.48), ranking above 34.22%. Patient N.P. also fell within the control range (*t*(9) = −0.37, *p* = .72, *z_cc_* = −0.39), ranking below 27.86%.

#### Weightlifting Task

This task assessed participants’ ability to infer object mass from human-object interactions. Participants viewed paired videos of the same actor or actress lifting visually identical containers of different weights and indicated which container appeared heavier. Critically, the inference here relied on kinematic information from observing other human agents acting on objects, instead of interactions between inanimate objects. Thus, the task could be solved based on interpreting perceived effort.

Control participants achieved a mean accuracy of 94.44% (*N* = 9, *SD* = 7.78%, range: 75.00% −100.00%). Patient I.A. scored 100.00%; Patient N.P. scored 96.88% (see Fig. 7). Modified t-tests showed that both patients performed within the control range: Patient I.A. (*t*(8) = 0.68, *p* = .52, *z_cc_* = 0.71), ranking above 48.29%; Patient N.P. (*t*(8) = 0.30, *p* = .77, *z_cc_* = 0.31), ranking above 22.56%.To assess systematic error patterns in controls, we conducted a repeated-measure ANOVA across weight combinations and found a significant effect of mass combination (*F*(1.99, 15.96) = 3.036, *p* = .08, η^2^_g_ = 0.13). However, post-hoc pairwise comparisons with Holm corrections did not reveal any significant difference among any pairs (all ps > 0.13), so no score adjustments were applied. Results indicate that both patients were able to interpret weight from the kinematics of other people’ actions.

#### Interim Summary & Discussion: Control Tasks

Overall, both I.A. and N.P. performed well in the control tasks. This was not surprising as I.A. performed at ceiling for the dynamic mass inference tasks, but it served as an important reference point for interpreting N.P.’s difficulties in judging the mass of objects from collisions and inter-object interactions in dynamic scenes. N.P. successfully discriminated different velocities of moving objects and inferred relative mass from human action contexts and static images of everyday scenarios, demonstrating intact knowledge of mass and how objects with different weights typically behave. Her poor performance in the collision paradigms therefore cannot be attributed to a general deficit in understanding mass as a concept or applying it in familiar contexts. Instead, she seems to have a hard time when making flexible mass inferences in dynamic object–object interactions, where simple heuristics are insufficient.

Inference of mass is only one aspect of what is broadly referred to as intuitive physical reasoning. In many real-world situations, we must also infer properties of dynamic scenes that are not intrinsic to a specific object, such as predicting when or where something will occur. While the literature on physical reasoning often groups these abilities together under the umbrella of naive physics (e.g., see Fischer et al., 2016; Mitko & Fischer, 2023; Schwettmann et al., 2019; Ullman et al., 2017), different types of inferences, such as estimating timing versus judging fixed physical properties like mass, may rely on distinct cues and cognitive mechanisms. To explore whether patients could reason about other aspects of dynamic events, we developed two exploratory tasks that focused specifically on the participants’ ability to make temporal predictions within a physical context (see Fig. 8).

**Figure 8.**
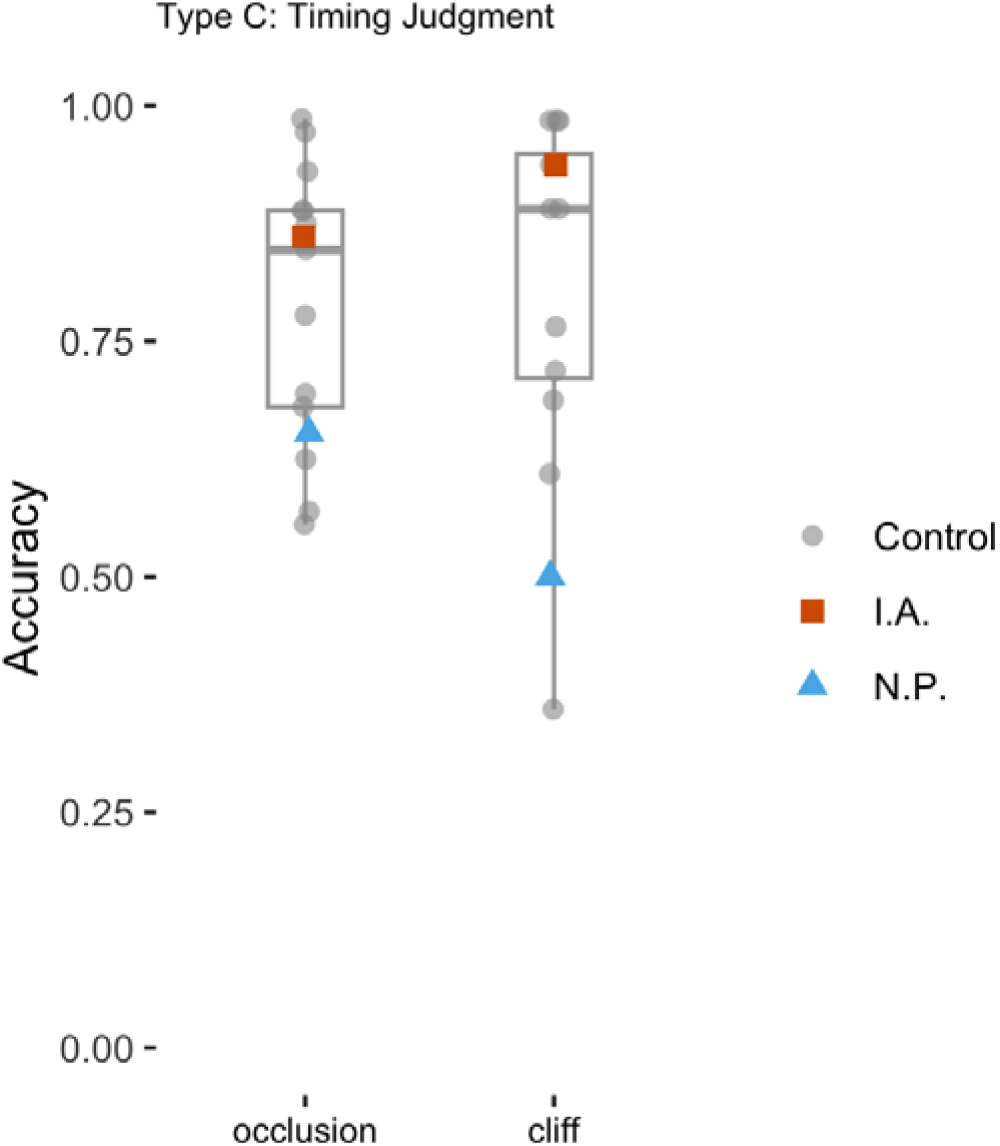
Accuracies for Type C - Timing Judgment tasks. The box plots showed the distribution of the control subjects. The individual dots showed the accuracy of individual subjects (gray circle: controls; red square: I.A.; blue triangle: N.P.).

### Type C: Timing tasks

#### Occlusion Task

This task probed participants’ ability to make predictions about object motion and timing. A ball rolled behind an occluder and reappeared either on time, early, or late. Participants judged whether the reappearance timing was correct. Control participants achieved a mean accuracy of 79.17% (*N* = 13, *SD* = 15.10%, range: 55.56% - 98.61%). Patient I.A. scored 86.11%; Patient N.P. scored 65.28%. There was no systematic error pattern across difficulty as indicated by difference from the expected timing (*F*(2, 24) = 1.77, *p* = .19, η^2^_g_ = 0.09), so no score adjustments were made. Both patients performed within the control range (I.A.: *t*(12) = 0.44, *p* = .67, *z_cc_* = 0.46, ranking above 33.45%; N.P.: *t*(12) = –0.89, *p* = .39, *z_cc_* = −0.92, ranking above 39.28%).

#### Cliff Timing Task

This task extended the occlusion paradigm by introducing vertical motion under gravity. Participants judged whether a falling ball reemerged from behind an occluder at the correct time. The correct timing depended solely on the ball’s initial horizontal velocity. Control participants performed with a mean accuracy of 81.25% (*N* = 12, *SD* = 19.19%, range: 35.94% - 98.44%). Patient I.A. scored 93.75%; Patient N.P. scored 50.00%. No systematic error pattern was found across timing deltas (*F*(2, 22) = 0.61, *p* = .55, η^2^_g_ = 0.02), so scores were not adjusted. Patient I.A.’s performance was within the control range (*t*(11) = 0.63, *p* = .54, *z_cc_* = 0.65), ranking above 45.57%. Patient N.P.’s performance was lower but not significantly so (*t*(11) = –1.56, *p* = .15, *z_cc_* = −1.63), ranking above 14.61%.

### Interim Summary & Discussion: Timing tasks

In the timing tasks, we assessed participants’ ability to make predictions about the temporal dynamics of a scene. While control participants exhibited a wide range of performance levels, I.A. achieved high accuracy in both tasks, despite their difficulty. Notably, N.P. also performed within the control range. This finding indicates that her difficulties in the collision paradigms cannot be attributed to a general inability to track motion or predict when objects will appear. In other words, although she struggled to infer weight from motion cues, she could accurately anticipate the timing of dynamic events, consistent with her ability to judge the velocity of objects with reasonable accuracy and demonstrating that her difficulty was not due to processing velocity or temporal information per se.

### General Discussion

Everyday behavior requires both reasoning about physical dynamics and using objects, yet whether these capacities rely on the same cognitive and neural mechanisms or can dissociate has remained an open question. In the present study, we examined physical reasoning and tool use in eleven left-hemisphere stroke patients. Tool use was assessed with a gesture-to-sight task, and physical reasoning (i.e., the ability to predict and interpret how objects react to external physical forces) with a novel set of nine tasks.

Within this sample, a key case illustrated a dissociation at the individual level: patient I.A., who despite being apraxic and impaired in tool-use pantomime, showed preserved physical reasoning and often outperformed neurotypical age-matched controls. This dissociation was complemented by patient N.P., who showed the opposite profile, with relatively preserved tool-use ability but marked difficulties in some physical reasoning tasks, particularly those requiring a flexible inference of mass based on dynamic interactions between pairs of objects. Furthermore, across the entire patient group, we found no systematic correlation between performance on our task battery (i.e., collision, timing, and control tasks) and praxis (see Appendix B, Table 2).

Taken together, these findings challenge the notion that tool use and physical reasoning rely on a common underlying system. Instead, they indicate that these two capacities, though deeply intertwined in everyday behavior, can be selectively disrupted by brain damage and likely depend on partially distinct cognitive and neural mechanisms. One might argue that the observed dissociation arises because the physical reasoning tasks we used were not directly tied to tool use. However, this interpretation actually reinforces our broader conclusion, namely, that both tool use and physical reasoning are multi-component constructs, and that claims of shared mechanisms should be made with caution when dealing with such complex cognitive domains.

### Physical reasoning and tool use are not unitary constructs

It has been proposed that physical reasoning, defined as “*the ability of reasoning about the physical properties of objects*”, broadly underlies action planning and tool use (Osiurak et al., 2024). The idea that physical reasoning and tool use rely on shared cognitive and neural mechanisms is, in part, motivated by the observation that the same frontoparietal regions have been consistently implicated both in the use of manipulable objects (Buxbaum & Saffran, 2002; Gallivan & Culham, 2015; Lewis, 2006; Reynaud et al., 2016) and in third-person physical reasoning tasks (Fischer et al., 2016; Schwettmann et al., 2019). This anatomical overlap has often been interpreted as evidence for shared mechanisms (Fischer & Mahon, 2021; Osiurak et al., 2024).

However, several assumptions in this interpretation remain under-tested. For instance, the neural bases of physical reasoning and tool use have rarely been examined within the same individuals, so the apparent overlap in brain activation remains largely untested within individuals. Moreover, because we interact with the physical world naturally and fluently, the underlying processes often co-occur, making them difficult to separate using neuroimaging or behavioral paradigms in neurotypical individuals alone. Lesion studies offer a complementary approach to studies done with neurotypical individuals as focal brain damage can lead to deficits in certain processes while leaving others intact.

Our findings complicate strong claims of a single shared mechanism underlying physical reasoning and tool use. While physical reasoning is likely important for many forms of object interaction, the present results challenge the idea that it is necessary or sufficient for all tool use (for a similar argument, see (Buxbaum, 2017). More broadly, our findings underscore the importance of distinguishing between different forms of physical reasoning and tool use, and how they relate to one another. Umbrella terms like *technical cognition* (Bartolo & Osiurak, 2022; Federico et al., 2025), *naive physics,* or object/tool use may obscure the heterogeneity in the cognitive and neural bases of these abilities and their different components.

We show that reasoning about object dynamics can be spared even when tool use is impaired (patient I.A.). Our data additionally suggest the opposite may also be true, namely that some aspects of physical reasoning can be impaired despite preserved ability to use tools (patient N.P.). Crucially, our findings extend this dissociation to third-person physical reasoning that do not involve direct interaction with objects, or reasoning about object affordances. Other ongoing work in our lab demonstrates that this dissociation also applies to reasoning about the ways that objects mechanically interact with each other more broadly (Du et al.).

We would like to note that even if future work were to show, with high precision, that physical reasoning and tool use recruit the same brain regions within individuals, such overlap would not imply that the two capacities are functionally identical or reducible to one another. Neural overlap across a set of regions may reflect partially shared representations, without each region contributing equally to both functions. Evidence from our patients illustrates this point. Patient I.A.’s lesion involved dorsal premotor and motor cortices, regions frequently identified as important for physical reasoning. Yet, he performed at ceiling across multiple tasks requiring the interpretation and prediction of object dynamics, despite struggling in tool use pantomime. This implies that an intact premotor cortex is not strictly necessary for third-person physical inference, at least in the perceptual judgment tasks used here. Note that Patient I.A.’s left supramarginal gyrus was intact. This region is widely recognized as a key hub for tool use (Bartolo & Osiurak, 2022; Federico et al., 2025; Osiurak et al., 2024) and has recently been proposed as the key cortical structure for the higher-order computations common to both physical reasoning and tool use (Fischer & Mahon, 2021; Navarro-Cebrián & Fischer, 2022). Yet, despite the integrity of this region, Patient I.A. was impaired in tool use while remaining unimpaired in third-person physical reasoning. This implies that this region on its own is not sufficient for tool use, although it does support the planning of movement trajectories, an ability known to be disrupted in limb apraxia (Wong et al. 2019). Perhaps more importantly, I.A.’s lesion disrupted the arcuate fasciculus, which is part of the ventro-dorsal stream of the praxis network that is critical for conveying information about tool actions from the temporal and parietal lobes to the motor system (Bi et al. 2015; Garcea et al. 2020; Howard et al. 2019).

Patient N.P., on the other hand, exhibited white matter lesions affecting dorsal fronto-parietal pathways, including the superior longitudinal fasciculus that connects the superior parietal lobule to dorsal premotor regions. The partial disconnection of this tract could account for the observed dissociation by selectively disrupting posterior–dorsal circuits involved in dynamic visuospatial integration and possibly in physical inference, while sparing the more inferior frontal and parietal regions connected via the arcuate fasciculus, that are associated with tool use and praxis. Together, these cases illustrate that even when lesions affect regions within the classical dorsal stream (i.e., the fronto-parietal system), the consequences for physical reasoning and tool use can diverge markedly based on the status of the ventro-dorsal versus dorso-dorsal divisions of that pathway (Rizzolatti and Matelli 2003; Binkofski and Buxbaum 2013). Both patients showed disruptions within dorsal pathways, implicating premotor and parietal regions often considered central to physical reasoning and tool use, yet their behavioral profiles were strikingly different. This dissociation underscores that similar anatomical loci can participate in distinct functional networks, and that successful tool use and physical reasoning rely on partially overlapping but dissociable mechanisms.

These observations are consistent with prior suggestions that it is the dorso-dorsal (rather than ventro-dorsal) pathway that is critical for mechanical reasoning about tool actions (Buxbaum 2017), and in line with the idea that brain regions often labeled as the *naive physics network* and *tool use network* can serve distinct functions in support of tool use or physical reasoning. For example, the posterior middle temporal gyrus has been linked to retrieving the visual appearance of tool-use actions, whereas the inferior parietal lobule (IPL) has been associated with object manipulation and with proprioceptive/somatosensory representations of how object manipulation feels (Buxbaum et al., 2007; Metzgar et al., 2022; Tobia & Madan, 2017). It has been proposed that within the IPL, the supramarginal gyrus may serve as a hub for integrating these aspects, buffering candidate tool-use actions before selection (Buxbaum et al., 2007; McDowell et al., 2018). However, the specific contributions of these regions to third-person physical reasoning remain unclear. Taken together, our findings and prior work caution against treating the “naive physics network” as a monolithic entity or assuming that its overlap with the tool-use network reflects a single, unified mechanism.Our task battery was designed to probe a range of inferences about fundamental physical properties of single objects such as mass and velocity. Some of these inferences were based on object-object interactions, others on human-object interactions; some relying on dynamic motion cues, others on static visual information. Although some studies propose that these different forms of reasoning draw on a shared cognitive resource (Mitko et al., 2024), our findings suggest a more differentiated picture. Inferring mass from dynamic collisions, reasoning from static cues, and predicting the timing of an event may overlap at some level, but they also require attention to and integration of distinct types of information. Our findings also suggest that not all forms of physical reasoning depend on a fully intact or flexible inferential system. Some tasks could be solved using simple heuristics, such as “*the heavier object moves more slowly*”, even if someone is not able to flexibly infer physical properties of an object from its dynamics and across contexts. This helps explain why N.P. succeeded on certain control tasks, such as estimating weight from the size of a splash, while struggling with collision tasks that required flexible integration of motion and force cues. Her performance illustrates that partial competence in physical reasoning can be supported by rules of thumb, without a fully abstract internal model or complete knowledge of the rules of physics.

Taken together, our findings highlight the importance of moving beyond umbrella terms like *naive physics* and *tool use*, and instead being specific about the types of representations and processes involved in different contexts. This approach can help us think more specifically about the types of processes that might be shared across the two capacities and which might dissociate. A more fine-grained approach is essential not only for understanding how these abilities are organized in the brain, but also for accurately characterizing the ways they break down in neurological populations.

### Future directions

Our study was based on behavioral profiles at the level of individual subjects and as such, it provides an existence proof for a dissociation between some aspects of tool use and physical reasoning. While identifying a dissociation between tool use and physical reasoning at the individual level provides evidence for their functional independence, studies with more diverse patient samples and experimental paradigms are needed to better characterize the scope and nature of this dissociation. More detailed investigations of individual patients will also be important for clarifying the nature of the observed deficits. For instance, although patient N.P. showed preserved tool use alongside difficulties in physical reasoning, are there specific components of tool use that might still be vulnerable? Similarly, for patient I.A., does his preserved physical reasoning abilities extend to reasoning about the physics of familiar tools, or are they restricted to more abstract or decontextualized physical events?

It is also important to note that our study focused on patients with left hemisphere damage, as such lesions are frequently associated with deficits in tool use and therefore provide a valuable opportunity to examine potential dissociations between tool use and physical reasoning. However, the dissociation we observed was demonstrated primarily at the behavioral level, and our data do not provide sufficient neural evidence to draw strong conclusions about the specific brain regions involved. In our ongoing work, we are combining detailed behavioral assessments with neuroimaging and lesion mapping approaches to better identify the neural substrates underlying these capacities.

There is growing interest in third-person physical reasoning and its connections to action planning, tool use, and other cognitive processes. While anatomical overlap between these domains has been noted in the literature, it is often based on findings from separate studies. Directly assessing different types of physical reasoning within the same individuals, both behaviorally and with neuroimaging, will be critical for clarifying the cognitive and neural bases of these capacities. For instance, future work on physical reasoning can distinguish between tasks that can be solved based on simple heuristics and those requiring more flexible, generalizable inference, as well as examine differences between reasoning in dynamic versus static contexts. As another example, paradigms can be designed in which tool-use interactions themselves require physical inference, for instance, adjusting one’s action as a function of an object’s weight. Some aspects of physical reasoning and tool use are likely to go hand in hand, and designing tasks that systematically vary the need for physical inference during action, or vice versa, will be key to identifying where their underlying mechanisms converge or dissociate.

## Conclusion

Our findings suggest that physical reasoning and tool use are both complex and heterogeneous processes that depend on dissociable systems. Although the brain regions supporting tool use and intuitive physics overlap anatomically to some extent, the cognitive processes they support can be selectively impaired. Claims about shared computations should be made with precision, carefully considering the types of reasoning involved, the task demands, and the brain regions in question.

## Acknowledgements

We are deeply grateful to the patients and control participants for their time and participation. We also thank John Yates for his assistance with data collection and Branch Coslett for his guidance in lesion segmentation. This work was supported by NIH Grant #R01NS115862 awarded to Aaron Wong and by European Research Executive Agency Widening programme under the European 5 Union’s Horizon Europe Grant 101087584 “CogBooster” awarded to Alfonso Caramazza.

## Appendix A

Scatterplots for praxis (measured by Gesture-to-Sight task) vs task performance. Control data is shown in a boxplot for comparison. *Legends: **Orange square** - patient I.A.; **Blue triangle** – patient N.P.; **Gray circles** – other patients*

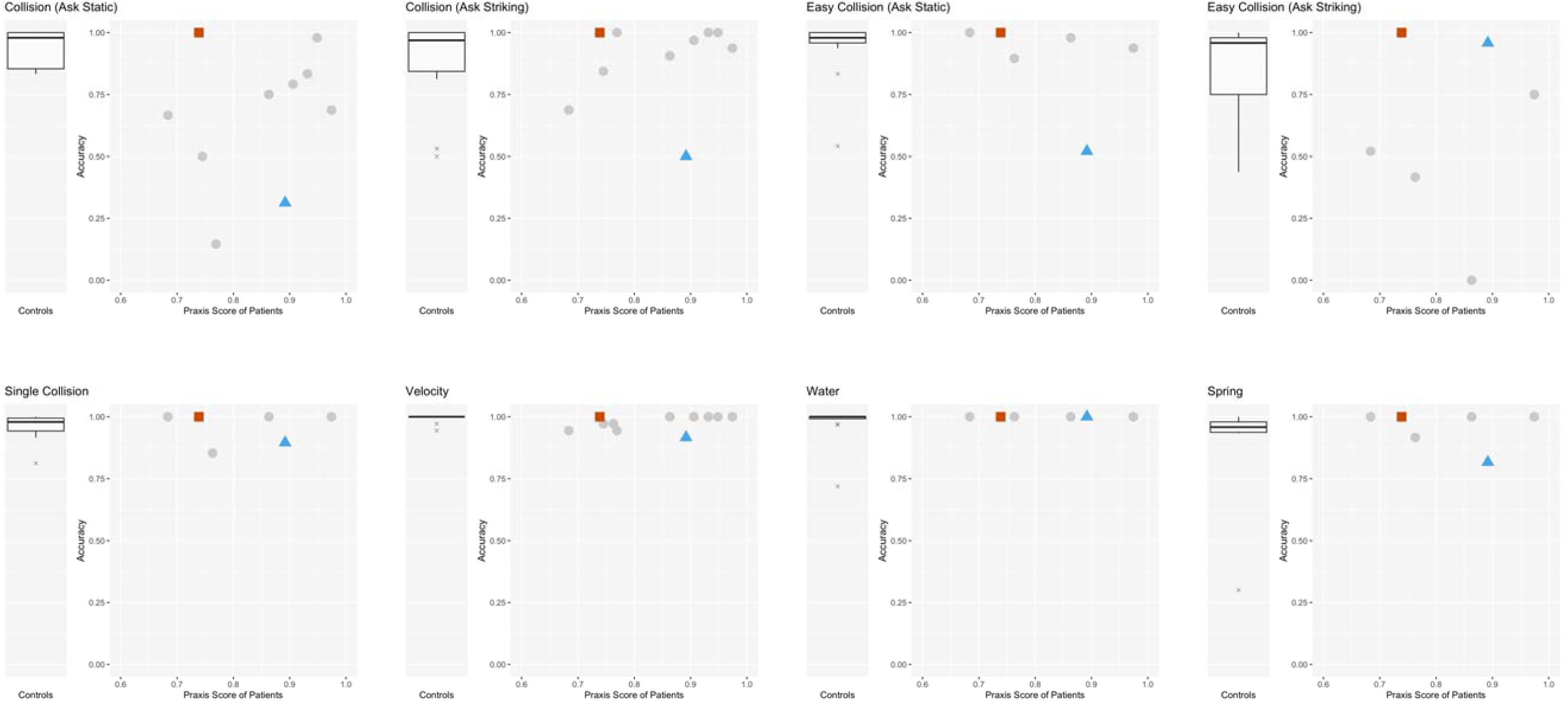
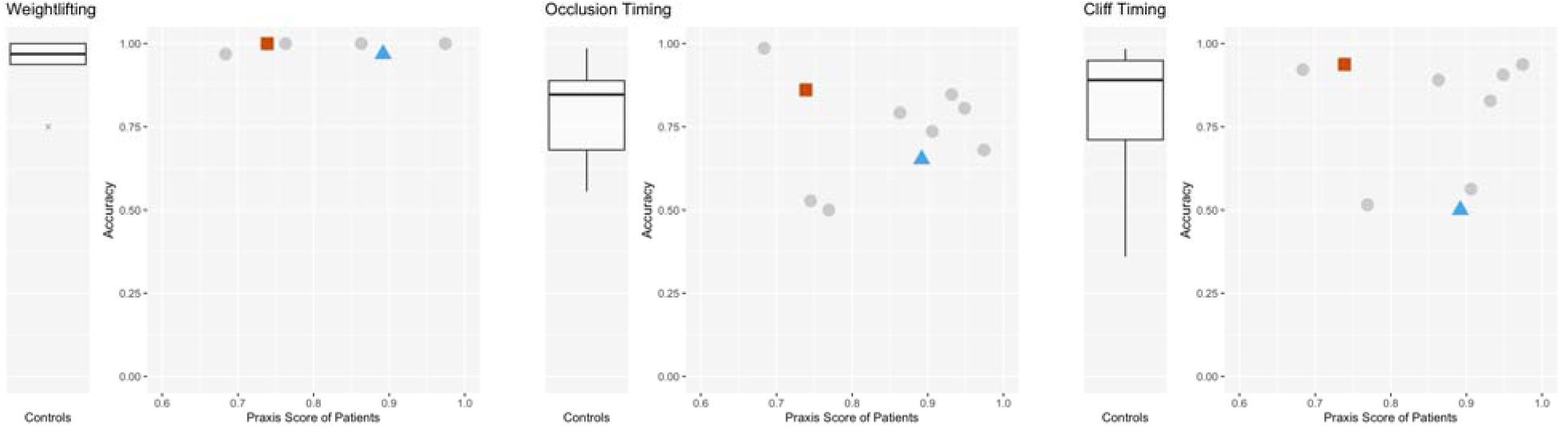

## Appendix B

**Table 1.** Task Accuracy of Control Group, Patient I.A., and Patient N.P.

| Task | Control Group |  |  |  |  | I.A. | N.P. |
| --- | --- | --- | --- | --- | --- | --- | --- |
|  | N | Mean | SD | Min | Max |  |  |
| Collision (Static) | 13 | 93.91% | 7.39% | 83.33% | 100.00% | 100.00% | 31.25% |
| Collision (Striking) | 13 | 80.93% | 22.37% | 37.50% | 100.00% | 100.00% | 52.08% |
| Adjusted Collision (Striking) | 13 | 89.18% | 17.80% | 50.00% | 100.00% | 100.00% | 50.00% |
| Easy Collision (Static) | 15 | 94.31% | 11.90% | 54.17% | 100.00% | 100.00% | 52.08% |
| Easy Collision (Striking) | 14 | 82.44% | 21.08% | 43.75% | 100.00% | 100.00% | 95.83% |
| Single Collision | 10 | 95.62% | 5.76% | 81.25% | 100.00% | 100.00% | 89.58% |
| Velocity | 13 | 93.11% | 8.35% | 75.00% | 100.00% | 100.00% | 70.83% |
| Adjusted Velocity | 13 | 99.36% | 1.66% | 94.44% | 100.00% | 100.00% | 91.66% |
| Water | 12 | 97.14% | 97.14% | 97.14% | 97.14% | 100.00% | 100.00% |
| Spring | 10 | 8.05% | 8.05% | 8.05% | 8.05% | 100.00% | 81.67% |
| Weightlifting | 9 | 94.44% | 7.78% | 75.00% | 100.00% | 100.00% | 96.88% |
| Occlusion | 13 | 79.17% | 15.10% | 55.56% | 98.61% | 86.11% | 65.28% |
| Cliff | 12 | 81.25% | 19.19% | 35.94% | 98.44% | 93.75% | 50.00% |

**Table 2.**
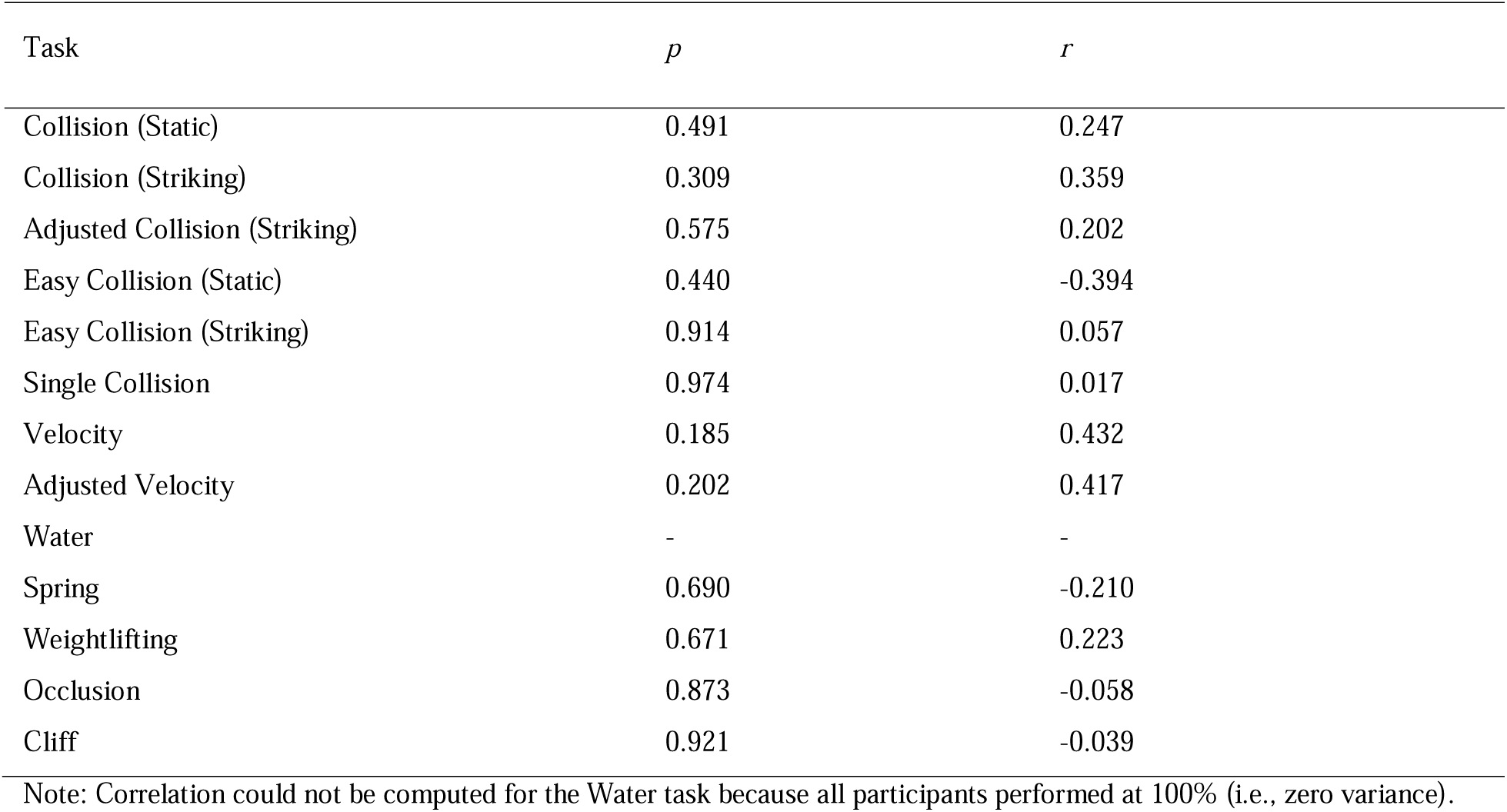
Pearson Correlation Between Task Accuracy And Praxis Score Of All Patients.

## Appendix C

White matter disconnection maps of patients I.A. and N.P. presented in MNI space, generated using the BCBToolkit (Foulon et al. 2018).

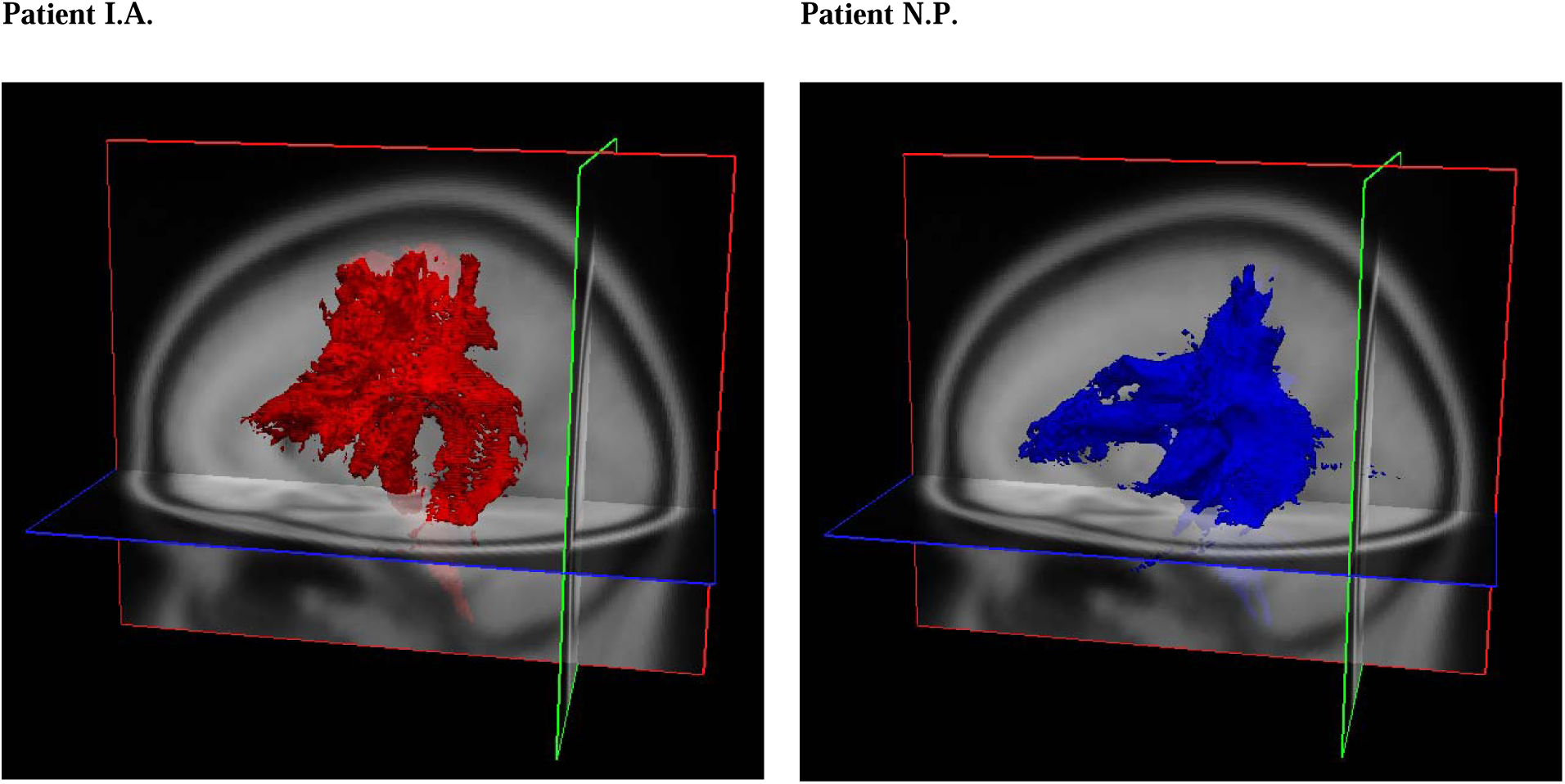

## Footnotes

* Both initials I.A. and N.P. are pseudonyms used to protect participant identities

